# Intrinsic excitability of rat hippocampal granule cells increases along the dorsal-to-ventral axis

**DOI:** 10.64898/2026.01.01.697271

**Authors:** Sanjna Kumari, Rishikesh Narayanan

**Affiliations:** Cellular Neurophysiology Laboratory, Molecular Biophysics Unit, Indian Institute of Science, Bengaluru 560012, India

**Keywords:** granule cells, hippocampus, heterogeneities, intrinsic excitability, patch-clamp electrophysiology

## Abstract

The dentate gyrus (DG) of the hippocampus exhibits striking anatomical and functional heterogeneity along its dorsoventral axis, yet the intrinsic electrophysiological diversity of its principal excitatory neurons, the granule cells, across the dorsoventral axis remains unexplored. Here, we systematically examined the electrophysiological properties of DG granule cells across the dorsal, intermediate, and ventral regions of the rat hippocampus. We found a progressive increase in input resistance, impedance amplitude, and firing rate of granule cells, accompanied by a gradual slowdown in repolarization kinetics of their action potentials along the dorsal-to-ventral axis. Our analyses demonstrated that granule cells acted as class I integrators that lacked strong resonance properties across the dorsoventral axis. We performed pairwise correlation and dimensionality reduction analyses to reveal weak dependencies across physiological measurements and the absence of distinct clusters for dorsal, intermediate, or ventral granule cells. Importantly, blade-specific analyses of granule cell physiology revealed that all measurements manifested pronounced heterogeneities even within a given dorsoventral section and a specific blade. Strikingly, ventral granule cells in the infrapyramidal blade manifested higher firing rates compared to their counterparts in the suprapyramidal blade. These blade-specific differences were limited to the ventral granule cells, with negligible distinctions between granule cells in the two blades of either dorsal or intermediate hippocampus. Together, our findings unveil a progressive increase in excitability of DG granule cells along the dorsal-to-ventral axis and a blade-specific granularity of firing properties, adding new dimensions to the several known anatomical, molecular, and behavioral differences across the hippocampal dorsoventral axis.

## INTRODUCTION

The hippocampus is functionally and anatomically organized along its dorsoventral (septotemporal) axis, with dorsal regions primarily implicated in spatial memory and navigation, and ventral regions contributing to emotion, motivation, and stress-related behaviors (Fanselow & Dong, 2010; Strange *et al*., 2014; Cembrowski *et al*., 2016; Cembrowski & Spruston, 2019). The dorsoventral functional dichotomy extends to differences in other attributes such as gene expression profiles (Cembrowski *et al*., 2016; Cembrowski & Spruston, 2019), afferent and efferent connectivity (Witter *et al*., 2000; Andersen *et al*., 2006; Witter, 2007), sensitivity to epileptogenic stimuli (Elul, 1964b, a; Racine *et al*., 1977; Gilbert *et al*., 1985; Bragdon *et al*., 1986; Arnold *et al*., 2019), and changes in adult neurogenesis in depression (Tanti & Belzung, 2013). At the cellular level, dorsoventral differences in intrinsic electrophysiological properties have been extensively characterized in CA1 pyramidal neurons (Dougherty *et al*., 2012; Marcelin *et al*., 2012; Hönigsperger *et al*., 2015; Kim & Johnston, 2015; Malik *et al*., 2016; Malik & Johnston, 2017) and in medial entorhinal cortex stellate cells (Giocomo *et al*., 2007; Garden *et al*., 2008; Giocomo & Hasselmo, 2008; Pastoll *et al*., 2020), suggesting that intrinsic excitability across the dorsoventral axis is regionally tuned to support distinct computational roles.

The dentate gyrus (DG) is the input gate to the hippocampus (Johnston & Amaral, 2004; Andersen *et al*., 2006; Amaral *et al*., 2007). The DG has been implicated in several cognitive behaviors including pattern separation (Marr, 1971; Andersen *et al*., 2006; Amaral *et al*., 2007; Morgan *et al*., 2007; Aimone *et al*., 2014; Borzello *et al*., 2023), novelty detection (Davis *et al*., 2004; Li *et al*., 2012; Kropff *et al*., 2015; Wu *et al*., 2015; Yang *et al*., 2017; Kesner, 2018; Hainmueller *et al*., 2024), spatial navigation (Danielson *et al*., 2017; GoodSmith *et al*., 2017; Senzai & Buzsaki, 2017; Zhang & Jonas, 2020; GoodSmith *et al*., 2022), and engram formation (Liu *et al*., 2012; Ramirez *et al*., 2013; Redondo *et al*., 2014; Ramirez *et al*., 2015; Ryan *et al*., 2015; Park *et al*., 2016; Roy *et al*., 2016; Stefanelli *et al*., 2016; Hainmueller & Bartos, 2018, 2020; Josselyn & Tonegawa, 2020; Sun *et al*., 2020) as well as in several pathological conditions such as epilepsy, Alzheimer’s disease, depression, and schizophrenia (Scharfman *et al*., 2000; Scharfman *et al*., 2002; Scharfman, 2007; Sutula & Dudek, 2007; Winner *et al*., 2011; Hagihara *et al*., 2013; Patricio *et al*., 2013; Yu *et al*., 2014; Cho *et al*., 2015; Leal & Yassa, 2015; Botterill *et al*., 2019). There are important differences in the behavioral and pathophysiological roles attributed to the dorsal *vs*. the ventral dentate gyrus (Pothuizen *et al*., 2004; Fanselow & Dong, 2010; Kheirbek *et al*., 2013; Weeden *et al*., 2014; Kempadoo *et al*., 2016; Kesner, 2018; Woods *et al*., 2020; Yeates *et al*., 2020; Dirven *et al*., 2022). For instance, the dorsal dentate gyrus has been implicated in exploratory drive, encoding of contextual fear memories, conjunctive encoding of multisensory inputs and spatial pattern separation. In contrast, the ventral dentate gyrus has been found to suppress innate anxiety while supporting odor pattern separation, encoding reward value, and social recognition (Kheirbek *et al*., 2013; Kempadoo *et al*., 2016; Kirk *et al*., 2017; Kesner, 2018).

Despite the critical role of the DG as the entry point to the hippocampal trisynaptic circuit, the question of whether granule cells (the principal cells of the DG) manifest dorsoventral gradients in their cellular properties remains unknown. Anatomically, within each section along the dorsoventral axis, the DG is further divided into the infrapyramidal and the suprapyramidal blades. Although there are several well-established differences across these two blades (Amaral *et al*., 2007), dorsoventral gradients in blade-specific differences in granule cell intrinsic physiology remain unexplored.

In this study, we systematically investigated the dorsoventral gradient of intrinsic excitability in granule cells of the dentate gyrus using whole-cell patch-clamp recordings from acute hippocampal slices. We report that ventral granule cells are significantly more excitable than their dorsal counterparts, displaying both enhanced subthreshold responsiveness and increased action potential firing. These differences formed a progressive dorsoventral gradient, with increasing excitability toward the ventral pole of the hippocampus. Blade-specific analyses revealed that ventral granule cells in the infrapyramidal blade manifested higher firing rates compared to those in the suprapyramidal blade, without significant differences in subthreshold excitability. Importantly, these blade-specific differences were limited to the ventral hippocampus and did not extend to dorsal or intermediate hippocampus. Together, our findings provide evidence for a hitherto unknown gradient in granule cell intrinsic excitability, highlighting both a progressive increase in excitability along the dorsal-to-ventral axis and a blade-specific diversity in firing properties. These findings suggest that dorsoventral gradients in granule cell excitability contribute to region-specific information processing in the hippocampus, potentially aligning granule cell physiology with the broader functional and anatomical organization of the hippocampal formation.

## MATERIALS AND METHODS

### Ethical approval

All experiments reported in this study were reviewed and approved by the Institute Animal Ethics Committee of the Indian Institute of Science, Bangalore. Experimental procedures were similar to previously established protocols (Narayanan & Johnston, 2007; Narayanan & Johnston, 2008; Ashhad *et al*., 2015; Ashhad & Narayanan, 2016; Das & Narayanan, 2017; Mishra & Narayanan, 2020a; Mishra & Narayanan, 2020b, 2022) and are detailed below. Experiments were performed on Sprague Dawley rats between 30–50 days of age. Animals were provided *ad libitum* food and water and were housed with an automated 12:12-h light-dark cycle, with the facility temperature maintained at 23° C. All animals were obtained from the in-house breeding setup at the Central Animal Facility of the Indian Institute of Science.

### Slice preparation for *in vitro* patch-clamp recording

Rats were anesthetized by intraperitoneal injection of a ketamine-xylazine mixture. After onset of deep anaesthesia, assessed by cessation of toe-pinch reflex, transcardial perfusion of ice-cold cutting solution was performed. The cutting solution contained (in mM) 2.5 KCl, 1.25 NaH_2_PO_4_, 25 NaHCO_3_, 0.5 CaCl_2_, 7 MgCl_2_, 7 dextrose, 3 sodium pyruvate, and 200 sucrose (pH 7.3, ∼300 mOsm) saturated with 95% O_2_-5% CO_2_. Thereafter, the brain was removed quickly and 350-µm thick slices were prepared from hippocampi with a vibrating blade microtome (Leica Vibratome) while submerged in ice-cold cutting solution saturated with 95% O_2_ - 5% CO_2_. For ventral and intermediate hippocampal slices, near horizontal sections were prepared as described in (Narayanan & Johnston, 2007; Narayanan & Johnston, 2008; Ashhad *et al*., 2015; Ashhad & Narayanan, 2016; Das & Narayanan, 2017; Mishra & Narayanan, 2020a; Mishra & Narayanan, 2020b, 2022). Specifically, these slices were prepared by laying either hemisphere of the brain on its sagittal surface. A blocking cut was made at around 30° relative to the vertical plane on the dorsal surface of the hemisphere. After that the tissue block was mounted on the flat surface created by the blocking procedure with the ventral region on the top of the block. Approximately 3 ventral slices and 3 intermediate slices were obtained from each hemisphere using this technique.

Dorsal hippocampal slices were prepared using established procedures (Dougherty *et al*., 2012; Malik *et al*., 2016), by first making a blocking cut at 45° angle from the coronal plane starting at the posterior end of the forebrain. A second blocking cut was also made at 45° relative to the coronal plane, but starting from approximately one-third of the total length of the forebrain from the most anterior point. Brains were mounted on the flat surface created by the first blocking cut. Approximately 4–5 dorsal hippocampal slices were obtained from each hemisphere. The near bregma and interaural coordinates for slices containing the hippocampus of different dorsoventral regions are as follows (Bregma, interaural in mm): dorsal (–4.74 to – 3.86, 5.2 to 6.14), intermediate (–5.6 to –6.1, 3.4 to 3.9), ventral (7.6 to –6.82, 2.4 to 3.18).

The slices were then incubated for 10–15 min at 34 °C in a chamber containing a holding solution (pH 7.3, ∼300 mOsm) with the composition of (in mM) 125 NaCl, 2.5 KCl, 1.25 NaH_2_PO_4_, 25 NaHCO_3_, 2 CaCl_2_, 2 MgCl_2_, 10 dextrose, and 3 sodium pyruvate saturated with 95% O_2_-5% CO_2_. Thereafter, the slices were kept in a holding chamber at room temperature for at least 30 min before the start of recordings.

### Electrophysiology: Whole cell current-clamp recording

For electrophysiological recordings, slices were transferred to the recording chamber and continuously perfused with carbogenated artificial cerebrospinal fluid (ACSF-extracellular recording solution) at a flow rate of 2–3 mL/min. All recordings were performed in current-clamp configuration at physiological temperatures (32–35 °C) achieved through an inline heater that was part of a closed-loop temperature control system (Harvard Apparatus). The carbogenated ACSF contained (in mM) 125 NaCl, 3 KCl, 1.25 NaH_2_PO_4_, 25 NaHCO_3_, 2 CaCl_2_, 1 MgCl_2_, and 10 dextrose (pH 7.3; ∼300 mOsm). Slices were first visualized under a 10× objective lens to focus on the infrapyramidal or suprapyramidal blades of the dentate gyrus (DG). Then a 63× water-immersion objective lens was employed to perform visually guided patch-clamp recordings from granule cells (GCs), through a contrast microscope (Carl Zeiss Axioexaminer). GCs were identified by the presence of their cell bodies within the granule cell layer and recordings were distributed to span the stretch of the infrapyramidal or suprapyramidal blades.

Whole cell current-clamp recordings were performed from neurons with Dagan BVC-700A amplifiers. Borosilicate glass electrodes with electrode tip resistance between 3 and 7 MΩ were pulled (P-97 Flaming/Brown micropipette puller; Sutter) from thick glass capillaries (1.5-mm outer diameter and 0.86-mm inner diameter; Sutter) and used for patch-clamp recordings. The pipette solution contained (in mM) 120 K-gluconate, 20 KCl, 10 HEPES, 4 NaCl, 4 Mg-ATP, 0.3 Na-GTP, and 7 K_2_-phosphocreatine (pH 7.3 adjusted with KOH; osmolarity ∼300 mOsm). Series resistance was monitored and compensated online with the bridge-balance circuit of the amplifier. Experiments were discarded only if the initial resting membrane potential was more depolarized than –60 mV and if series resistance rose above 50 MΩ or if there were fluctuations in temperature and ACSF flow rate during the course of the experiment. Voltages have not been corrected for the liquid junction potential, which was experimentally measured to be ∼8 mV.

### Pharmacological blockers

All recordings were performed in the presence of synaptic receptor blockers in the ACSF. Drugs and their concentrations used in the experiments were 10 µM 6-cyano-7-nitroquinoxaline-2,3-dione (CNQX), an AMPA receptor blocker; 10 µM (+) bicuculline and 10 µM picrotoxin, both GABA_A_ receptor blockers; and 2 µM CGP55845, a GABA_B_ receptor blocker in the bath solution. All synaptic blockers were procured from Abcam.

### Subthreshold measurements

We characterized neuronal subthreshold properties with several established electrophysiological protocols (Narayanan & Johnston, 2007; Narayanan & Johnston, 2008; Ashhad *et al*., 2015; Ashhad & Narayanan, 2016; Das & Narayanan, 2017; Mishra & Narayanan, 2020a; Mishra & Narayanan, 2020b, 2022). Input resistance (*R*_*in*_) was measured as the slope of a linear fit to the steady-state voltage response *vs*. current amplitude (*V* − *I*) plot obtained by injecting subthreshold current pulses of amplitudes spanning –25 to +25 pA, in steps of 5 pA. To assess temporal summation, five-alpha excitatory postsynaptic potentials (*⍺*-EPSPs) with 50-ms interval were evoked by current injections of the form *I*_α_(*t*) = *I*_max_ *t* exp(– α*t*), with *⍺* = 0.1 ms^−1^. Temporal summation ratio (*S*_*⍺*_) in this train of five EPSPs was computed as *E*_*last*_/*E*_*first*_, where *E*_*last*_and *E*_*first*_are the amplitudes of the last and first EPSPs in the train, respectively. Sag ratio (*Sag* in %) was measured from the voltage response of the cell to a hyperpolarizing current pulse of 100 pA and was defined as 100 × 71 − *V*_*SS*_/*V*_*peak*_8, where *V*_*ss*_ and *V*_*peak*_ depict the steady-state and peak voltage deflection from the resting membrane potential (*V*_*RMP*_), respectively.

The chirp stimulus used for characterizing the impedance amplitude (|*Z*|) and phase (Φ) profiles was a sinusoidal current of constant amplitude designed to yield voltage responses below firing threshold, with its frequency linearly spanning 0–15 Hz in 15 s (referred to as the *Chirp15* stimulus). A 50-pA hyperpolarizing current pulse was provided before the chirp current to estimate input resistance as well as to observe and correct series resistance changes through the course of the experiment. The voltage response of the neuron to the *Chirp15* current stimulus was recorded at *V*_*RMP*_. The magnitude of the ratio of the Fourier transform of the voltage response to the Fourier transform of the *Chirp15* stimulus formed the impedance amplitude profile (ZAP). The associated phase formed the impedance phase profile (ZPP). The frequency at which the impedance amplitude reached its maximum (|*Z*|_max_) was the resonance frequency (*f*_*R*_). Resonance strength (*Q*) was measured as the ratio of the maximum impedance amplitude to the impedance amplitude at 0.5 Hz. Total inductive phase (Φ_L_) was defined as the area under the inductive (positive values) part of the ZPP (Narayanan & Johnston, 2008). Unless otherwise stated, all subthreshold measurements were measured at *V*_RMP_. In a subset of experiments, negative or positive currents were injected to alter the membrane potential (in the range –85 to –60 mV in steps of 5 mV) to measure subthreshold properties (*R*_*in*_, *Sag*, *f*_*R*_, |*Z*|_max_, *Q*, and Φ_L_) at different membrane voltages.

### Suprathreshold measurements

Action potential (AP) firing frequency was computed by extrapolating the number of spikes obtained during a 700-ms current injection to 1 s. Current amplitude of these pulse current injections was varied from 0 pA to 250 pA in steps of 50 pA to access the lower current injection regime and 500 pA to 1250 pA in steps of 250 pA to access higher current injection regime. The notation *f*_*X*_was used to present the firing rate of the neuron to an *X* pA current injection. These values were used to construct the firing frequency *vs*. injected current (*f*–I) plots for low and high current injection ranges. Various AP-related measurements (Mishra & Narayanan, 2020a) were derived from the voltage response of the cell to a 250-pA pulse current injection. AP amplitude (*V*_*AP*_) was computed as the difference between the peak voltage of the spike (*V*_peak_) and *V*_RMP_. The temporal distance between the timing of the first spike and the time of current injection was defined as the latency to first spike (*T*_1AP_). The duration between the first and the second spikes was measured as the first inter-spike interval (*T*_1ISI_). AP half-width (*T*_APHW_) was the temporal width measured at the half-maximal points of the AP peak with reference to *V*_RMP_. The maximum (*dV*/*dt*|_max_) and minimum (*dV*/*dt*|_min_) rate of change of voltage of the AP temporal derivative were calculated from the temporal first derivative of the AP trace. The voltage in the AP trace corresponding to the time point at which the *dV*/*dt* crossed 20 V/s defined AP threshold (*V*_*th*_). All suprathreshold measurements were obtained through current injections into the cell resting at *V*_RMP_.

### Data acquisition and analyses

All data acquisition and analyses were performed with custom-written software in IGOR Pro 6 (WaveMetrics). To avoid misinterpretations arising from representation of merely the summary statistics and to emphasize heterogeneities within and across groups, all individual data points are explicitly represented. Statistical analyses were performed using non-parametric tests. The Kruskal-Wallis test was used to determine whether there are significant differences between three or more independent groups. The Wilcoxon rank sum was used to assess significant differences between two groups. All statistical analyses were performed using the R computing package (http://www.r-project.org/).

## RESULTS

We performed whole-cell patch-clamp electrophysiological recordings in current-clamp mode at the cell body of visually identified granule cells in the three different sections of hippocampus: dorsal, intermediate and ventral. We compared subthreshold and suprathreshold properties of granule cells from these different sections as well as between the infrapyramidal and suprapyramidal blades of the dentate gyrus.

### Subthreshold excitability of dentate gyrus granule cells increased along the dorsal-to-ventral axis

We characterized subthreshold electrophysiological properties of granule cells of the dentate gyrus across the dorsoventral axis of the hippocampus (Figs. 1–2, Tables 1–2). Specifically, we measured input resistance (Fig. 1*B–C*), temporal summation (Fig. 1*D*), sag ratio (Fig. 1*E*) and frequency-dependent impedance measurements (Figs. 1*F–H*) from granule cells across the dorsoventral axis. We found that granule cells showed resting membrane potentials (RMP) in the range of –80 to –65 mV with no significant differences across the different sections (Fig. 2*A*; Tables 1–2). However, input resistance (a subthreshold measure of neuronal excitability) manifested a progressive increase along the dorsal-to-ventral axis, with significantly lower values in dorsal GCs compared to their intermediate and ventral counterparts (Fig. 2*B*; Tables 1–2). Temporal summation was greater than unity across most recorded granule cells and did not show significant differences across the dorsoventral axis (Fig. 2*C*; Tables 1–2). The sag was consistently low (<10 %) and was not significantly different across the dorsal-to-ventral axis (Fig. 2*D*; Tables 1–2).

**Figure 1.**
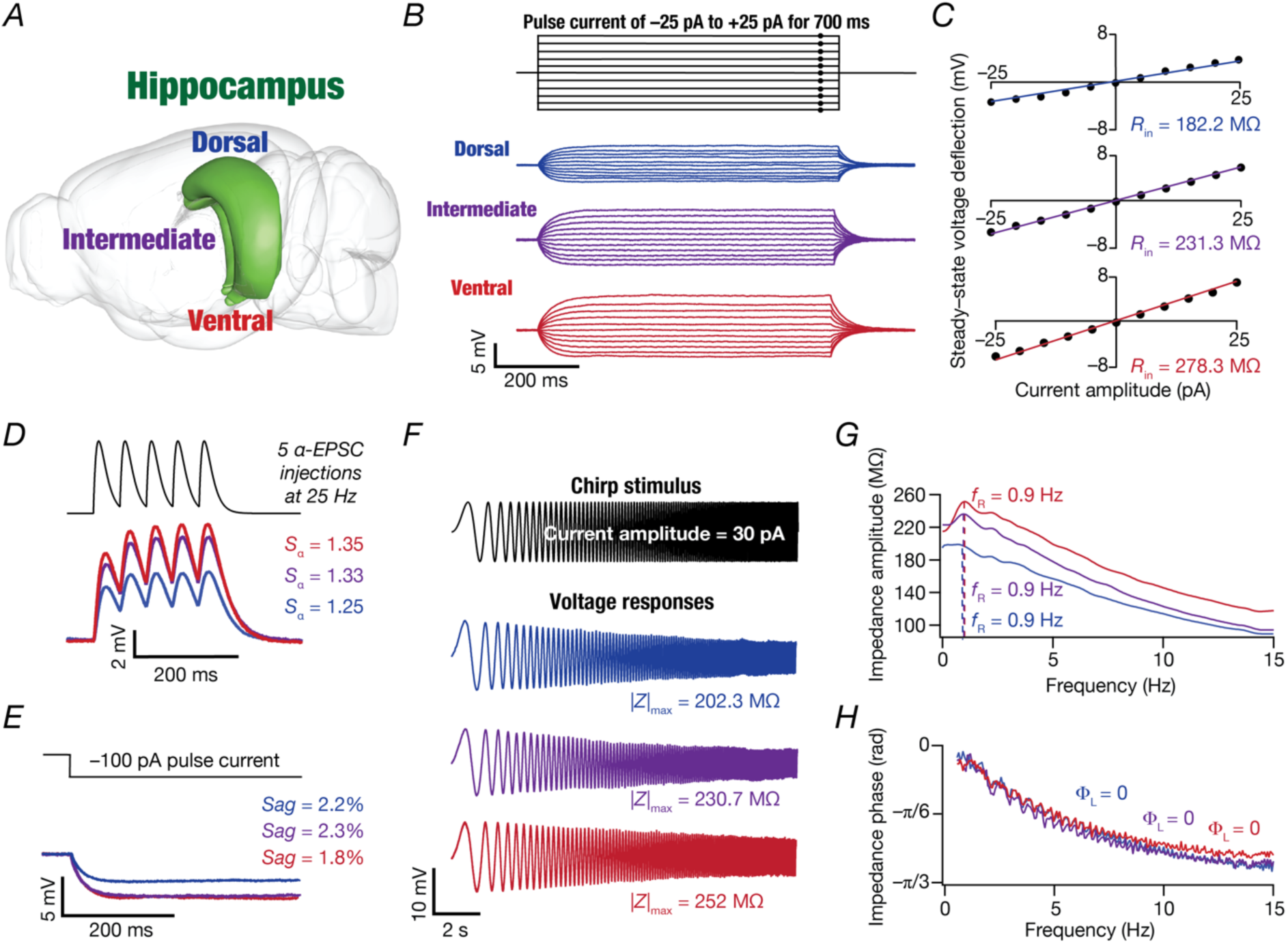
Illustrative example of electrophysiological characterization of subthreshold excitability of dentate gyrus granule cells across the dorsoventral axis. *A,* Schematic of rodent brain showing the anatomical structure of dorsal, intermediate, and ventral hippocampal subregions (Image from Scalable Brain Atlas (Bakker *et al*., 2015)). *B*, Voltage responses of example neurons in each of the dorsal, intermediate and ventral subregions to 700-ms current pulses of amplitude varying from –25 pA to +25 pA in steps of 5 pA shown on top. *C*, *V* − *I* plots derived from the steady-state voltage deflections obtained from the example traces shown in panel B. Input resistance (*R*_*in*_) was calculated as the slope of the plot depicting steady-state voltage response as a function of the injected current amplitude. *D*, Somatic voltage response to five alpha excitatory postsynaptic current (α-EPSC) injections at 25 Hz for all the three example neurons to calculate temporal summation ratio (*S*_*⍺*_). *S*_*⍺*_ was computed as the ratio of the amplitude of the fifth response to that of the first. *E*, Somatic voltage response from all the three example neurons to a 100 pA hyperpolarizing current injection. Sag was computed from the steady-state (*V*_*SS*_) and peak (*V*_*peak*_) values of voltage deflections from resting membrane potential (*V*_*RMP*_). *F*, *Top*: Chirp current stimulus spanning 0–15 Hz in 15 s of 30 pA amplitude. *Bottom*: Somatic voltage response to chirp current injection from example cells from dorsal, intermediate, and ventral GCs. *G–H*, Impedance amplitude (G) and phase (H) profiles plotted as a function of frequency obtained from the chirp current stimulus for all the three example neurons. *All example traces are derived from the same representative neuron, one each from dorsal, intermediate, and ventral regions*.

**Figure 2.**
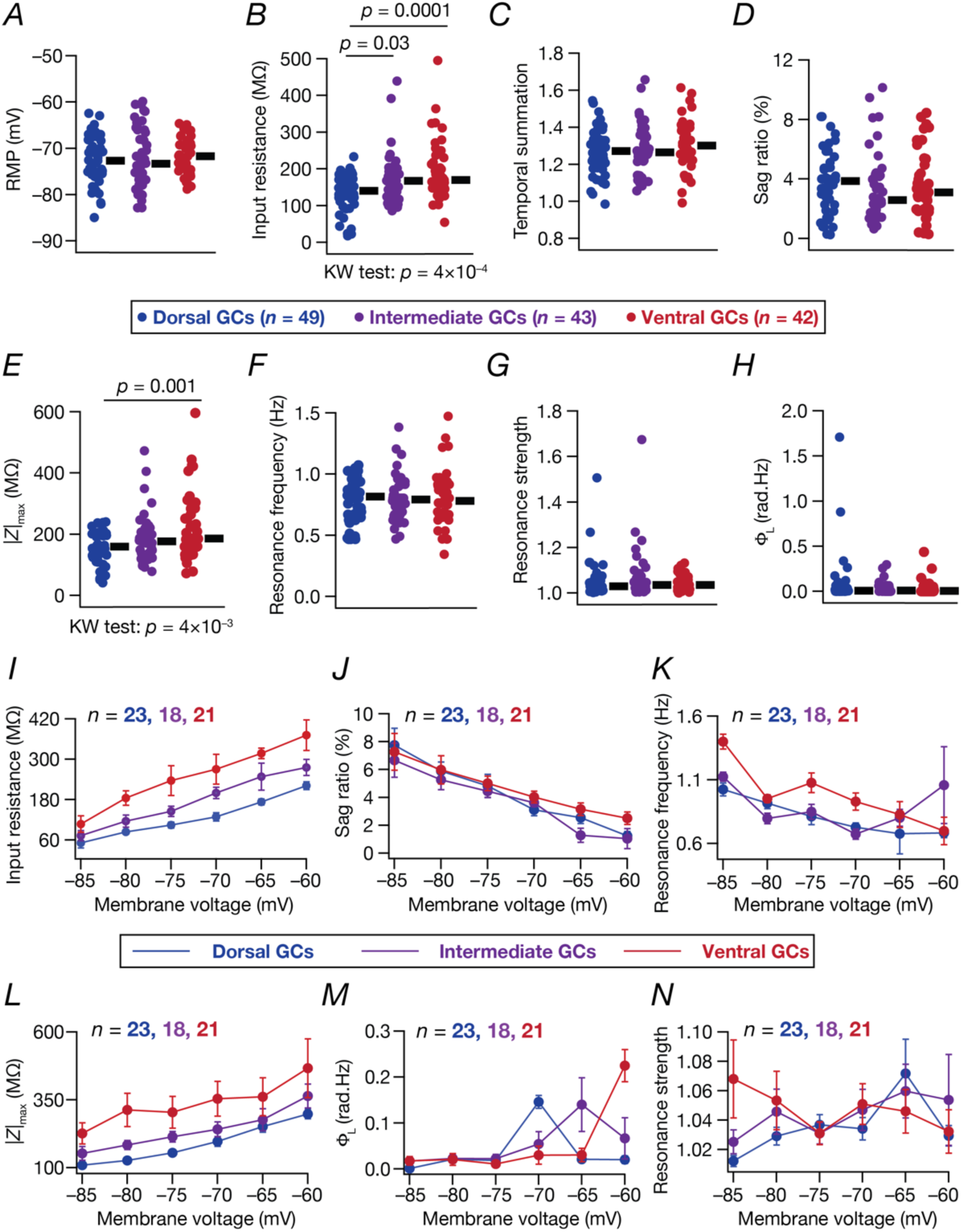
Subthreshold excitability of hippocampal granule cells increased along the dorsoventral axis of the dentate gyrus. *A–H*, Beeswarm plots depicting subthreshold measurements from granule cells of dorsal, intermediate, and ventral dentate gyrus: (A) resting membrane potential, *V*_*RMP*_(mV); (B) Input resistance, *R*_*in*_(MΩ); (C) Temporal summation, *S*_*⍺*_; (D) Sag (%); (E) Maximal impedance amplitude, |*Z*|_*max*_ (MΩ); (F) Resonance frequency, *f*_*R*_ (Hz); (G) Resonance strength, *Q*; and (H) Total inductive phase, Φ_L_(rad.Hz) from all the recorded granule cells from dorsal (*n* = 49), intermediate (*n* = 43) and ventral (*n* = 42) dentate gyrus. The thick lines on right of each beeswarm plot represent the median for the specified population. All measurements depicted in this figure were obtained through current injections into the neuron resting at resting membrane potential, *V*_*RMP*_. *I–N*, Voltage dependence of steady-state (I and J) and frequency-dependent (K–N) measurements of GCs across the dorsoventral axis of granule cells in dorsal (*n* = 23), intermediate (*n* = 18), and ventral (*n* = 21) granule cells. Data is represented with mean and SEM. *Statistical test: Significant p values obtained from the Kruskal–Wallis (KW) test are mentioned below the graphs. Wilcoxon rank sum test was performed between two groups, significant p values are mentioned on the top of figures. Additional summary statistics are provided in Table 1. All statistical test outcomes associated with this figure are provided in Tables 2–3*.

**Table 1:** Summary statistics for electrophysiological measurements recorded from neurons in dorsal, intermediate and ventral GCs when respective current protocols were injected with cell resting at *V*_RMP_ (Fig. 2*A–H*, Fig. 4*B–J*). Measurements are mean ± SEM (*n* cells) and the degree of variability in each measurement is reported as standard deviation, interquartile range, and coefficient of variation, in that order.

| Measurement, unit | Symbol | Dorsal GCs | Intermediate GCs | Ventral GCs |
| --- | --- | --- | --- | --- |
| *Subthreshold measurements* |  |  |  |  |
| Resting membrane potential, mV | $V_{RMP}$ | $-72.8 \pm 0.7$ (49)<br>5.12, 6.09, -0.07 | $-71.9 \pm 1.0$ (42)<br>6.8, 11.5, -0.09 | $-71.7 \pm 0.6$ (43)<br>3.79, 5.28, -0.05 |
| Input resistance, MΩ | $R_{in}$ | $134.5 \pm 7.1$ (49)<br>49.6, 53.5, 0.36 | $171 \pm 11.2$ (42)<br>71.93, 69.23, 0.42 | $195.8 \pm 12.2$ (43)<br>80.3, 77.3, 0.41 |
| Temporal summation ratio | $S_{\alpha}$ | $1.2 \pm 0.01$ (49)<br>0.12, 0.12, 0.09 | $1.27 \pm 0.02$ (42)<br>0.14, 0.21, 0.11 | $1.31 \pm 0.02$ (43)<br>0.14, 0.16, 0.01 |
| Sag ratio, % | $Sag$ | $3.78 \pm 0.3$ (47)<br>2.09, 2.66, 0.55 | $3.36 \pm 0.36$ (42)<br>2.36, 2.63, 0.70 | $3.59 \pm 0.33$ (43)<br>2.18, 2.68, 0.61 |
| Resonance frequency, Hz | $f_R$ | $0.78 \pm 0.02$ (47)<br>0.16, 0.2, 0.2 | $0.79 \pm 0.02$ (42)<br>0.19, 0.24, 0.23 | $0.80 \pm 0.03$ (43)<br>0.22, 0.24, 0.28 |
| Maximum impedance amplitude, MΩ | $\vert Z\vert _{max}$ | $156.6 \pm 7.6$ (47)<br>52.2, 71.1, 0.33 | $189 \pm 12.3$ (42)<br>79.8, 80.3, 0.4 | $228.3 \pm 16.6$ (43)<br>108.8, 127.93, 0.47 |
| Resonance strength | $Q$ | $1.05 \pm 0.01$ (47)<br>0.08, 0.03, 0.07 | $1.07 \pm 0.02$ (42)<br>0.11, 0.05, 0.11 | $1.04 \pm 0.005$ (43)<br>0.03, 0.03, 2.2 |
| Total inductive phase, rad·Hz | $\Phi_L$ | $0.08 \pm 0.04$ (47)<br>0.27, 0.03, 3.1 | $0.03 \pm 0.01$ (42)<br>0.06, 0.04, 1.8 | $0.03 \pm 0.01$ (43)<br>0.07, 0.03, 2.2 |
| *Suprathreshold measurements* |  |  |  |  |
| Firing rate at 50 pA, Hz | $f_{50}$ | 0(49) | $0.1 \pm 0.1$ (41)<br>0.7, 0, 5.02 | $1.1 \pm 0.5$ (41)<br>3.33, 0, 3.08 |
| Firing rate at 100 pA, Hz | $f_{100}$ | $0.69 \pm 0.17$ (49)<br>1.2, 1.4, 1.7 | $1.4 \pm 0.4$ (41)<br>2.88, 1.42, 2.06 | $3.9 \pm 1.2$ (41)<br>7.89, 4.28, 1.98 |
| Firing rate at 150 pA, Hz | $f_{150}$ | $3.58 \pm 0.48$ (49)<br>3.3, 5.7, 0.94 | $6.41 \pm 0.9$ (41)<br>5.82, 7.14, 0.9 | $9.98 \pm 1.64$ (41)<br>10.5, 10, 1.05 |
| Firing rate at 200 pA, Hz | $f_{200}$ | $7.52 \pm 0.8$ (49)<br>5.6, 10, 0.75 | $12.9 \pm 1.5$ (41)<br>9.8, 11.42, 0.75 | $15.45 \pm 1.94$ (41)<br>12.4, 14.28, 0.08 |
| Firing rate at 250 pA, Hz | $f_{250}$ | $12.5 \pm 1.0$ (49)<br>7.06, 10, 0.56 | $19.2 \pm 2.1$ (42)<br>13.5, 11.42, 0.70 | $20.7 \pm 2.05$ (42)<br>13.3, 18.2, 0.64 |
| Firing rate at 500 pA, Hz | $f_{500}$ | $30.7 \pm 2.8$ (43)<br>18.7, 27.14, 0.6 | $50.8 \pm 4.4$ (40)<br>27.8, 29.07, 0.54 | $43.8 \pm 4.2$ (36)<br>25.09, 30.35, 0.57 |
| Firing rate at 750 pA, Hz | $f_{750}$ | $41.02 \pm 3.98$ (42)<br>25.85, 37.14, 0.63 | $66.1 \pm 5.0$ (39)<br>31.24, 46.42, 0.47 | $55.05 \pm 5.13$ (35)<br>30.37, 35.71, 0.55 |
| Firing rate at 1000 pA, Hz | $f_{1000}$ | $35.14 \pm 4.2$ (42)<br>27.5, 40, 0.78 | $59.4 \pm 5.8$ (39)<br>36.21, 50.71, 0.61 | $38.51 \pm 4.19$ (35)<br>24.83, 39.28, 0.64 |
| Firing rate at 1250 pA, Hz | $f_{1250}$ | $23.6 \pm 3.4$ (42)<br>20.15, 37.5, 0.85 | $41.8 \pm 5.2$ (39)<br>32.23, 44.3, 0.77 | $24.0 \pm 3.4$ (34)<br>20.07, 29.64, 0.83 |
| Peak action potential voltage, mV | $V_{peak}$ | $40.4 \pm 1.3$ (45)<br>8.96, 11.5, 0.22 | $42.7 \pm 1.6$ (40)<br>10, 14.5, 0.23 | $42.4 \pm 1.6$ (41)<br>10.5, 10.2, 0.24 |
| Action potential amplitude, mV | $V_{AP}$ | $112.3 \pm 1.41$ (45)<br>9.4, 12.25, 0.084 | $114.7 \pm 1.2$ (40)<br>7.86, 8.3, 0.06 | $113.3 \pm 1.6$ (41)<br>10.46, 13.14, 0.09 |
| Action potential threshold, mV | $V_{th}$ | $-37.6 \pm 1.1$ (45)<br>7.4, 10.4, -0.19 | $-34.4 \pm 1.2$ (40)<br>7.09, 8.36, -0.2 | $-33.1 \pm 1.2$ (41)<br>8.3, 12.3, -0.25 |
| Action potential halfwidth, ms | $T_{APHW}$ | $0.86 \pm 1.03$ (45)<br>0.25, 0.2, 0.28 | $1.01 \pm 0.03$ (40)<br>0.2, 0.27, 0.21 | $1.04 \pm 0.04$ (41)<br>0.29, 0.35, 0.27 |
| Peak dV/dt, V/s | $dV/dt\vert _{max}$ | $409.8 \pm 13.9$ (45)<br>92.9, 139.7, 0.22 | $357.5 \pm 14$ (40)<br>94.4, 121.9, 0.26 | $358.7 \pm 20.3$ (41)<br>130.34, 219.1, 0.36 |
| Minimum dV/dt, V/s | $dV/dt\vert _{min}$ | $-121.2 \pm 4.1$ (45)<br>27.6, 28.1, -0.22 | $-110.3 \pm 4$ (40)<br>25.3, 21.2, -2.22 | $-104.5 \pm 4.5$ (41)<br>28.6, 34.8, -0.27 |
| Latency to first spike, ms | $T_{1AP}$ | $29.08 \pm 2.76$ (45)<br>18.53, 23.2, 0.63 | $40.1 \pm 5.22$ (40)<br>33.05, 31.9, 0.82 | $42.87 \pm 9.88$ (41)<br>63.3, 29.4, 1.5 |
| First inter-spike interval, ms | $T_{1ISI}$ | $23.35 \pm 7.96$ (45)<br>53.4, 14.4, 2.28 | $22.2 \pm 5.73$ (40)<br>35.32, 14.54, 1.58 | $20.90 \pm 5.98$ (39)<br>37.3, 7.81, 1.78 |

**Table 2:** Statistical test results for data shown in (Fig. 2*A–H*, Fig. 4*B–J*) performed between different groups (D: Dorsal, I: Intermediate, V: Ventral). KW: Kruskal-Wallis test. Statistically significant *p* values are represented in boldface font.

| Measurement, unit | Symbol | KW test | Pairwise Wilcoxon Rank sum test |  |  |
| --- | --- | --- | --- | --- | --- |
|  |  |  | *V* vs. *I* | V vs. D | I vs. D |
| *Subthreshold measurements* |  |  |  |  |  |
| Resting membrane potential, mV | $V_{\text{RMP}}$ | 0.56 | 0.58 | 0.25 | 0.75 |
| Input resistance, MΩ | $R_{in}$ | **0.0004** | 0.09 | **0.0001** | **0.03** |
| Temporal summation ratio | $S_{\alpha}$ | 0.259 | 0.18 | 0.12 | 0.98 |
| Sag ratio, % | $Sag$ | 0.351 | 0.36 | 0.44 | 0.18 |
| Resonance frequency, Hz | $f_R$ | 0.913 | 0.93 | 0.8 | 0.6 |
| Maximum impedance amplitude, MΩ | $\vert Z\vert _{max}$ | **0.004** | 0.09 | **0.001** | 0.12 |
| Resonance strength | $Q$ | 0.843 | 0.78 | 0.77 | 0.55 |
| Total inductive phase, rad·Hz | $\Phi_L$ | 0.806 | 0.82 | 0.46 | 0.80 |
| *Suprathreshold measurements* |  |  |  |  |  |
| Firing rate at 50 pA, Hz | $f_{50}$ | **0.014** | 0.13 | **0.006** | 0.12 |
| Firing rate at 100 pA, Hz | $f_{100}$ | 0.144 | 0.11 | 0.08 | 0.8 |
| Firing rate at 150 pA, Hz | $f_{150}$ | **0.0005** | 0.07 | **0.0002** | **0.01** |
| Firing rate at 200 pA, Hz | $f_{200}$ | **0.0008** | 0.40 | **0.0006** | **0.004** |
| Firing rate at 250 pA, Hz | $f_{250}$ | **0.0017** | 0.51 | **0.001** | **0.006** |
| Firing rate at 500 pA, Hz | $f_{500}$ | **0.0010** | 0.22 | **0.01** | **0.0003** |
| Firing rate at 750 pA, Hz | $f_{750}$ | **0.0015** | 0.19 | **0.039** | **0.0003** |
| Firing rate at 1000 pA, Hz | $f_{1000}$ | **0.0035** | **0.008** | 0.42 | **0.0023** |
| Firing rate at 1250 pA, Hz | $f_{1250}$ | **0.0067** | **0.01** | 0.78 | **0.004** |
| Peak action potential voltage, mV | $V_{\text{peak}}$ | 0.5478 | 0.93 | 0.33 | 0.36 |
| Action potential amplitude, mV | $V_{AP}$ | 0.4828 | 0.88 | 0.45 | 0.20 |
| Action potential threshold, mV | $V_{th}$ | **0.0479** | 0.51 | **0.02** | 0.07 |
| Action potential halfwidth, ms | $T_{\text{APHW}}$ | **0.0005** | 0.95 | **0.001** | **0.0006** |
| Peak dV/dt, V/s | $dV/dt\vert _{\text{max}}$ | 0.0643 | 0.87 | 0.11 | **0.01** |
| Minimum dV/dt, V/s | $dV/dt\vert _{\text{min}}$ | **0.0032** | 0.38 | **0.002** | **0.007** |
| Latency to first spike, ms | $T_{1\text{AP}}$ | 0.2168 | 0.40 | 0.45 | 0.07 |
| First inter-spike interval, ms | $T_{1\text{ISI}}$ | 0.3208 | 0.08 | 0.51 | 0.586 |

**Table 3:** Statistical analyses associated with voltage dependence of subthreshold measurements (Fig. 2*I–N*). Statistically significant *p* values are represented in boldface font.

| Measurement,<br>unit | Dorsal |  | Intermediate |  | Ventral |  | KW test | Pairwise Wilcoxon Rank sum test |  |  |
| --- | --- | --- | --- | --- | --- | --- | --- | --- | --- | --- |
|  | Mean | SEM | Mean | SEM | Mean | SEM |  | *V* vs. *I* | *V* vs. *D* | *I* vs. *D* |
| Membrane potential –60 mV |  |  |  |  |  |  |  |  |  |  |
| Input resistance, MΩ | 221.5 | 11.6 | 275.4 | 23.6 | 371.4 | 45.4 | 0.49 | 0.88 | **0.04** | 0.98 |
| Sag ratio, % | 1.24 | 0.31 | 1.04 | 0.72 | 2.5 | 0.45 | 0.13 | 0.93 | 0.05 | 0.11 |
| Resonance frequency, Hz | 0.68 | 0.04 | 1.06 | 0.3 | 0.7 | 0.11 | 0.14 | 0.16 | 0.07 | 0.62 |
| Maximum impedance amplitude, MΩ | 298.2 | 17.6 | 364.2 | 43.6 | 466.3 | 108.2 | 0.56 | 0.76 | 0.37 | 0.37 |
| Resonance strength | 1.03 | 0.01 | 1.05 | 0.03 | 1.03 | 0.01 | 0.07 | 0.22 | 0.03 | 0.23 |
| Total inductive phase, rad·Hz | 0.02 | 0.01 | 0.07 | 0.04 | 0.23 | 0.04 | 0.16 | 0.89 | 0.1 | 0.08 |
| Membrane potential –65 mV |  |  |  |  |  |  |  |  |  |  |
| Input resistance, MΩ | 173.4 | 8.7 | 248.1 | 40.3 | 317.4 | 15.8 | **0.006** | 0.55 | **0.005** | 0.37 |
| Sag ratio (%) | 2.54 | 0.42 | 1.28 | 0.51 | 3.16 | 0.45 | 0.51 | 0.77 | 0.38 | 0.29 |
| Resonance frequency, Hz | 0.68 | 0.16 | 0.8 | 0.07 | 0.83 | 0.1 | 0.6 | 0.47 | 0.34 | 0.88 |
| Maximum impedance amplitude, MΩ | 251.6 | 20.9 | 277.3 | 40.3 | 361.1 | 69.7 | 0.6 | 0.48 | 0.33 | 0.99 |
| Resonance strength | 1.07 | 0.02 | 1.06 | 0.02 | 1.05 | 0.01 | 0.88 | 0.9 | 0.79 | 0.57 |
| Total inductive phase, rad·Hz | 0.02 | 0.01 | 0.14 | 0.06 | 0.03 | 0.01 | 0.53 | 0.28 | 0.44 | 0.81 |
| Membrane potential –70 mV |  |  |  |  |  |  |  |  |  |  |
| Input resistance, MΩ | 129.0 | 12.3 | 199.7 | 17.1 | 270.3 | 44.1 | **0.002** | 0.29 | **0.001** | **0.04** |
| Sag ratio (%) | 3.1 | 0.42 | 3.62 | 0.29 | 4.03 | 0.41 | 0.13 | 0.54 | 0.07 | 0.15 |
| Resonance frequency, Hz | 0.73 | 0.03 | 0.67 | 0.04 | 0.93 | 0.07 | 0.8 | 0.62 | 0.84 | 0.54 |
| Maximum impedance amplitude, MΩ | 196.7 | 18.34 | 241.6 | 27.1 | 354.1 | 64.09 | **0.02** | 0.18 | **0.01** | 0.07 |
| Resonance strength | 1.03 | 0.01 | 1.05 | 0.01 | 1.05 | 0.01 | 0.9 | 0.92 | 0.71 | 0.72 |
| Total inductive phase, rad·Hz | 0.15 | 0.01 | 0.05 | 0.03 | 0.03 | 0.02 | 0.35 | 0.19 | 0.23 | 1 |
| Membrane potential –75 mV |  |  |  |  |  |  |  |  |  |  |
| Input resistance, MΩ | 104.3 | 8.98 | 145 | 15.2 | 236.4 | 44.1 | **0.0007** | **0.006** | **0.0001** | 0.08 |
| Sag ratio (%) | 4.82 | 0.83 | 4.44 | 0.45 | 5.01 | 0.53 | 0.66 | 0.47 | 0.41 | 0.91 |
| Resonance frequency, Hz | 0.81 | 0.06 | 0.85 | 0.06 | 1.08 | 0.08 | **0.004** | **0.002** | **0.02** | 0.31 |
| Maximum impedance amplitude, MΩ | 155.4 | 8.98 | 214.5 | 19.2 | 304.3 | 58.1 | **0.008** | 0.38 | **0.007** | **0.01** |
| Resonance strength | 1.04 | 0.01 | 1.03 | 0.01 | 1.03 | 0.01 | 0.79 | 0.52 | 0.66 | 0.76 |
| Total inductive phase, rad·Hz | 0.02 | 0.01 | 0.02 | 0.01 | 0.01 | 0.01 | 0.41 | 0.58 | 0.14 | 0.67 |

| **Membrane potential –80 mV** |  |  |  |  |  |  |  |  |  |  |
| --- | --- | --- | --- | --- | --- | --- | --- | --- | --- | --- |
| Input resistance, MΩ | 84.3 | 5.8 | 115.9 | 18.3 | 184.9 | 21.6 | **0.0001** | **0.008** | **0.0001** | 0.36 |
| Sag ratio (%) | 5.89 | 0.64 | 5.27 | 0.71 | 5.98 | 1.01 | 0.38 | 0.6 | 0.49 | 0.16 |
| Resonance frequency, Hz | 0.92 | 0.04 | 0.8 | 0.04 | 0.95 | 0.03 | 0.35 | 0.98 | 0.25 | 0.19 |
| Maximum impedance amplitude, MΩ | 127.1 | 8.8 | 183.6 | 16.3 | 313.2 | 61.6 | **0.0009** | 0.11 | **0.0008** | **0.007** |
| Resonance strength | 1.03 | 0.01 | 1.05 | 0.02 | 1.05 | 0.02 | 0.15 | 0.38 | 0.05 | 0.29 |
| Total inductive phase, rad·Hz | 0.02 | 0.01 | 0.02 | 0.01 | 0.02 | 0.01 | 0.39 | 0.19 | 0.58 | 0.4 |
| **Membrane potential –85 mV** |  |  |  |  |  |  |  |  |  |  |
| Input resistance, MΩ | 51.4 | 15.2 | 72.9 | 15.2 | 107.4 | 24.6 | 0.38 | 0.74 | 0.12 | 0.59 |
| Sag ratio (%) | 7.74 | 1.21 | 6.67 | 1.22 | 7.27 | 1.32 | 0.41 | 0.68 | 0.32 | 0.32 |
| Resonance frequency, Hz | 1.03 | 0.05 | 1.12 | 0.04 | 1.4 | 0.06 | 0.39 | 0.46 | 0.71 | 0.17 |
| Maximum impedance amplitude, MΩ | 110.0 | 11.2 | 152.9 | 25.2 | 226.3 | 39.2 | 0.09 | 0.21 | 0.08 | 0.17 |
| Resonance strength | 1.01 | 0.0003 | 1.03 | 0.01 | 1.07 | 0.03 | 0.9 | 0.68 | 0.93 | 0.93 |
| Total inductive phase, rad·Hz | 0.0007 | 0.0007 | 0.02 | 0.01 | 0.02 | 0.01 | 0.71 | 0.67 | 0.46 | 0.93 |

The impedance amplitude (*e.g.*, Fig. 1*G*) and phase (*e.g.*, Fig. 1*H*) profiles for granule cells across the dorsal-to-ventral axis showed low-pass, integrator-like characteristics lacking strong resonance characteristics. Quantitatively, we found a significant and progressive increase in the maximal impedance amplitude, a frequency-dependent measure of neuronal excitability, across the dorsal-to-ventral axis (Fig. 2*E*; Tables 1–2). Impedance measurements across the dorsoventral granule cells confirmed the integrator-like characteristics with resonance frequency consistently lower than 1 Hz for most cells (Fig. 2*F*; Tables 1–2), resonance strength hovering just above unity (Fig. 2*G*; Tables 1–2), and small values of total inductive phase (Fig. 2*H*; Tables 1–2). Importantly, all subthreshold measurements showed widespread heterogeneities (quantified in Table 1 using measures of variability) within each section (Fig. 2*A–H*).

These subthreshold measurements (Fig. 2*A–H*) were computed at resting membrane potential of the respective cells. Therefore, we asked if the gradients observed in input resistance and maximal impedance amplitude would extend to other voltages as well. To address this, we recorded neuronal responses at different voltages to compute input resistance (Fig. 2*I*), sag ratio (Fig. 2*J*), and impedance-related properties (Fig. 2*K–N*). We found input resistance (Fig. 2*I*; Table 3) and maximum impedance amplitude (Fig. 2*L*; Table 3) to significantly reduce and sag to significantly increase with hyperpolarization (Fig. 2*J*; Table 3). The three other measurements — resonance frequency (Fig. 2*K*; Table 3), total inductive phase (Fig. 2*M*; Table 3), and resonance strength (Fig. 2*N*; Table 3) — did not show strong trends as functions of membrane voltage. Importantly, the dorsoventral gradients in input resistance (Fig. 2*I*; Table 3) and maximum impedance amplitude (Fig. 2*L*; Table 3) were observed across all measured voltage ranges.

Together, heterogeneous subthreshold measurements from granule cells revealed a progressive gradient in subthreshold excitability measures (input resistance and maximal impedance amplitude) along the dorsal-to-ventral axis of the hippocampus. These excitability measurements were highest in ventral cells, lowest in dorsal cells, whereas intermediate cells showed measurements between the dorsal and ventral extremes. Our measurements also demonstrated that granule cells acted as integrators and lacked strong resonance properties across the dorsoventral axis.

### Action potential firing frequency of dentate gyrus granule cells increased along the dorsal-to-ventral axis

Do dorsoventral gradients in subthreshold excitability properties translate to gradients in action potential firing properties and other suprathreshold electrophysiological properties? To answer this, we injected depolarizing current pulses of different amplitudes to assess the firing profiles of granule cells across the dorsoventral axis (Fig. 3*A*; Tables 1–2). In addition, we quantified several metrics associated with action potentials (Fig. 3*B–C*; Tables 1–2) of DG granule cells across the dorsoventral axis.

**Figure 3:**
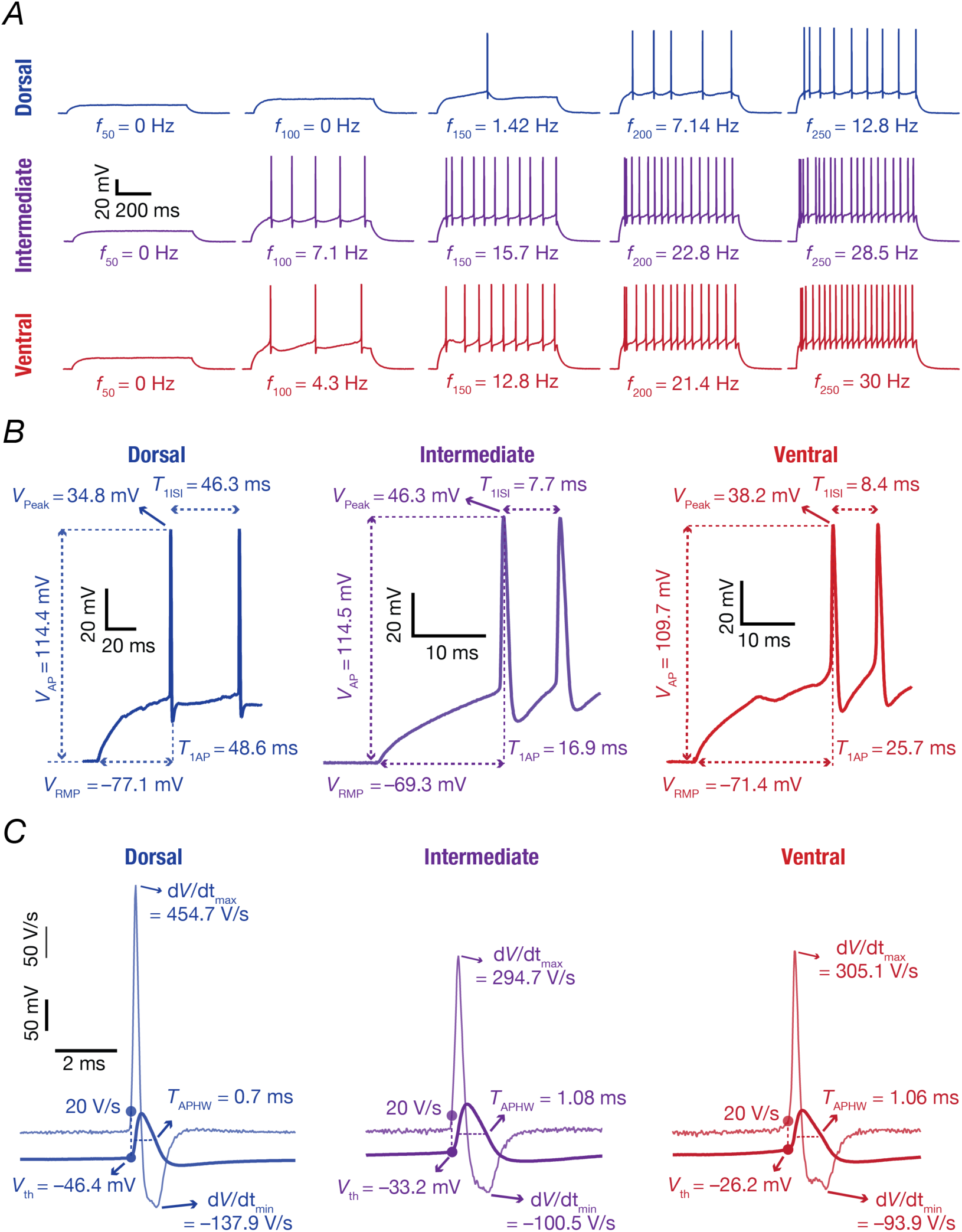
Illustrative example of electrophysiological characterization of suprathreshold excitability of dentate gyrus granule cells across the dorsoventral axis. *A*, Voltage responses of the example granule cells from dorsal, intermediate, and ventral dentate gyrus to a 700-ms current pulses from 50 pA to 250 pA in steps of 50 pA. *B,* Zoomed version of the trace shown in panel A for 250 pA current injection, showing the first two action potentials to illustrate the calculation of different action potential (AP) measurements. *V*_*RMP*_, resting membrane potential; *T*_1*AP*_, latency to the first action potential, measured from the time where the current injection was initiated; *V*_*peak*_, the maximum voltage value measured on the first action potential; *V*_*AP*_, the action potential amplitude, measured as the difference between *V*_*peak*_ and *V*_*RMP*_; *T*_1*ISI*_, first interspike interval measured as the temporal distance between the first and the second action potentials. *C*, Further zoomed version of the trace in panel A showing only the first action potential (thick traces), along with its temporal derivative (*dV*/*dt*, thin traces) illustrating the peak value of the derivate, *dV*/*dt*|_*max*_ and minimum value *dV*/*dt*|_*min*_. The voltage value at the timepoint at which temporal derivative crossed 20 V/s was assigned as the action potential threshold voltage (*V*_*th*_). The width at half-maximum of the action potential amplitude was measured as *T*_*APHW*_. *All example traces are derived from the same representative neurons, one each from dorsal, intermediate, and ventral regions (Same as* Fig. 1).

Across all three DG sections, consistent with integrator-like impedance profiles, we found the firing profile of granule cells (Fig. 3*A*, Fig. 4*B*, Tables 1–2) to exhibit class I excitability characteristics, with the ability to elicit action potentials at arbitrarily low frequencies (Hodgkin, 1948; Ratte *et al*., 2013; Mishra & Narayanan, 2020a). Strikingly, suprathreshold excitability was highest for ventral granule cells compared to their intermediate and dorsal counterparts (Fig. 4*B*, Tables 1–2). We inferred this from the ability of ventral granule cells to elicit significantly more number of action potentials for lower current injections (Fig. 4*B*; left, Tables 1–2) and by the early transitions to depolarization-induced block (DIB) for higher current injections (Fig. 4*B*; right, Tables 1–2). Specifically, a highly excitable neuron reaches high-firing rates that are unsustainable with the rate of sodium-channel recovery at relatively lower current values compared to neurons endowed with lower excitability. Thus, DIB-induced reduction in action potential firing frequencies for such highly excitable neurons occur at relatively lower current amplitudes. Consistent with the high excitability of ventral granule cells, it may be observed that the ventral population transitions from high-to-low firing frequencies at relatively low current values (Fig. 4*B*). Together, the dorsal-to-ventral increase in subthreshold excitability measurements (Fig. 2) extended to suprathreshold excitability as well (Fig. 4*B*).

**Figure 4.**
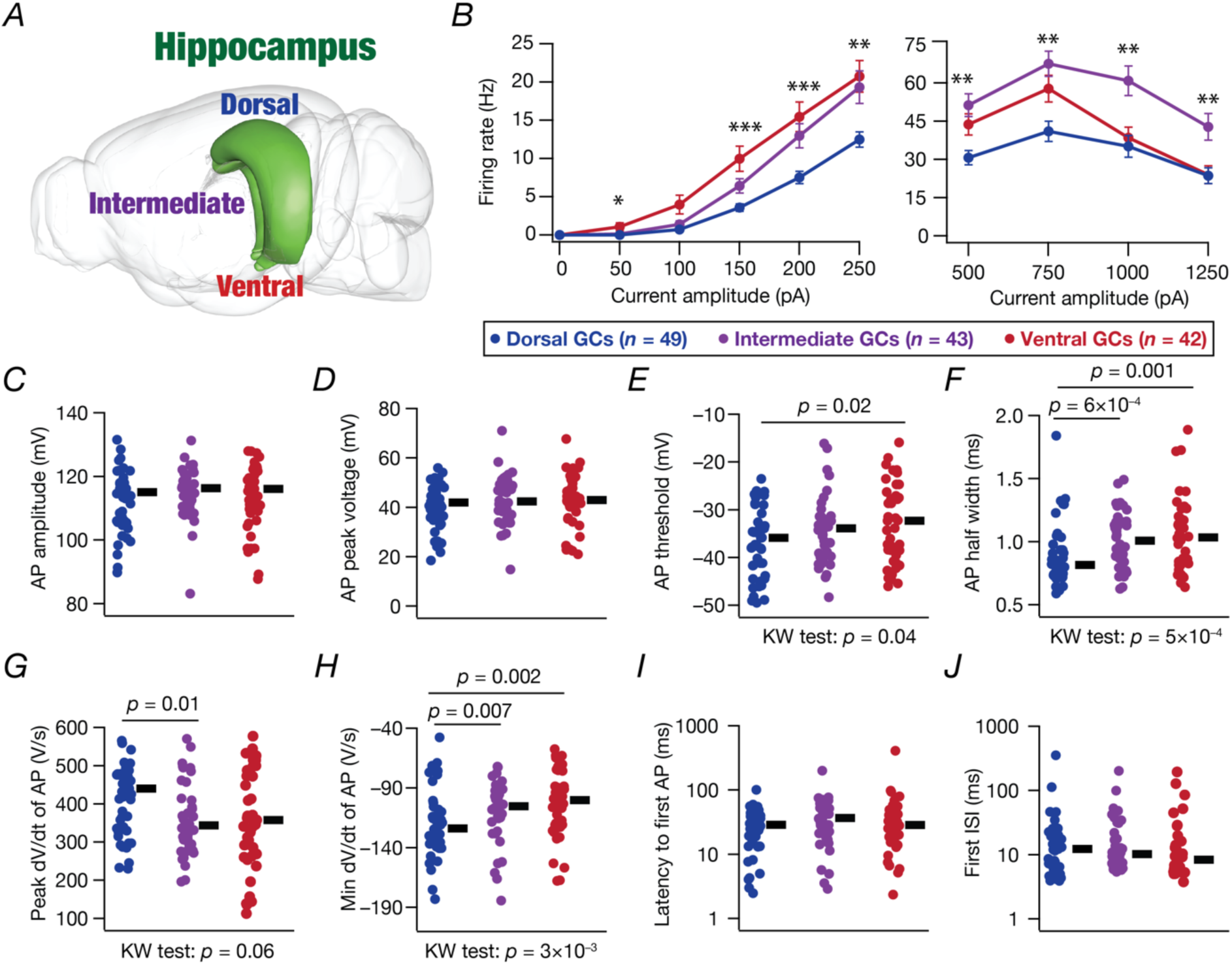
Suprathreshold excitability of hippocampal granule cells increased along the dorsoventral axis of the dentate gyrus. *A*, Schematic of rodent brain showing the anatomical structure of dorsal, intermediate, and ventral hippocampus (Image from Scalable Brain Atlas (Bakker *et al*., 2015)). *B, f* − *I* curve of action potential firing rate against injected current amplitude, ranging from 0 to 250 pA in steps of 50 pA (left) and for higher current injection from 500 pA to 1250 pA in steps of 250 pA (right). *p values: * <0.05, ** < 0.005, *** < 0.0005*. *C–J*, Beeswarm plots for action potential (AP) measurements calculated from the respective voltage responses obtained for the 250 pA pulse-current injection (as illustrated in Fig. 3). *Statistical test: Significant p values obtained from the Kruskal–Wallis (KW) test are mentioned below the graphs. Wilcoxon rank sum test was performed between two groups, significant p values are mentioned on the top of figures. Additional summary statistics are provided in Table 1. All statistical test outcomes associated with this figure are provided in Table 2*.

Turning to action potential (AP) measurements (Fig. 3*B–C*, Fig. 4*C–J*), we found significant dorsoventral differences in AP threshold (Fig. 4*E*, Tables 1–2), half width (Fig. 4*F*, Tables 1–2), and the peak as well as the minimum values of the first temporal derivative of action potential trace (Fig. 4*G–H*, Tables 1–2). On the other hand, AP amplitude (Fig. 4*C*, Tables 1–2), AP peak voltage (Fig. 4*D*, Tables 1–2), latency to first AP (Fig. 4*I*, Tables 1–2), and first inter-spike interval (Fig. 4*J*, Tables 1–2) did not show significant changes across granule cells of the dorsal-to-ventral axis. Action potential amplitude was largely above 100 mV (Fig. 4*C*). Although AP threshold showed significant differences between the dorsal and ventral populations, the difference in the mean/median was 3–4 mV, a relatively small shift compared to the heterogenous range of the values spanning around 30 mV (Fig. 4*E*, Tables 1–2). Similarly, the shift in peak *dV*/*dt* should be considered relatively small albeit being significant between the dorsal and intermediate populations, especially considering the variability (Fig. 4*G*, Tables 1–2).

The strongest dorsoventral gradients and the most significant differences in AP properties were associated with the two measures related to AP repolarization: AP half width (Fig. 4*F*, Tables 1–2) and minimum *dV*/*dt* of the AP (Fig. 4*H*, Tables 1–2). Specifically, there was a progressive increase in both AP half width and min *dV*/*dt* along the dorsal-to-ventral axis. A lower value of min *dV*/*dt* (Fig. 4*H*) implies a faster repolarization and would translate to a lower value of AP half width (Fig. 4*F*), together making these two observations consistent with each other. These observations together indicate a progressive dorsal-to-ventral gradient in AP repolarization kinetics, with dorsal neurons manifesting the thinnest action potentials (Fig. 4*F*) and the fastest repolarization (Fig. 4*H*).

Together, heterogeneous suprathreshold measurements from granule cells revealed a progressive gradient in firing rate and repolarization kinetics of action potentials along the dorsal-to-ventral axis of the hippocampus. Action potential firing rates were the highest in ventral granule cells and lowest in dorsal granule cells. Action potential repolarization kinetics were the fastest in dorsal granule cells and slowest in ventral granule cells. Our measurements also demonstrated that granule cells acted as class I integrators across the dorsoventral axis.

### A large proportion of the electrophysiological measurements across the dorsoventral axis of the dentate gyrus exhibited weak pairwise correlations

We next sought to understand the correlations between the different subthreshold and suprathreshold measurements. Are there strong relationships between them? Were the different subthreshold and suprathreshold measurements characterizing distinct aspects of granule cell physiology or did they quantify similar physiological characteristics? Was the measurements space high-dimensional or low-dimensional? Do dorsal, intermediate, and ventral neurons show distinct clusters in a reduced-dimensional space? To address these questions, we first computed pairwise correlations between the different measurements we obtained from neurons across all sections, where the complete set of measurements were available (*n* = 117). We plotted pairwise scatterplots of all intrinsic measurements and computed Pearson’s correlation coefficient for each of these pairwise scatterplots (Fig. 5*A*). We found weak pairwise correlations, ranging between –0.5 and 0.5, between most pairs (Fig. 5*A*). Specifically, there was a strong positive correlation between *R*_*in*_and |*Z*|_*max*_, both of which are measures of subthreshold excitability. There were also strong positive correlations between firing rates computed for different current injections. A large subset of uncorrelated measurements suggest that the set of measurements employed here in characterizing DG granule cells are assessing distinct aspects of their physiology. Among AP properties, there were strong positive correlations between *V*_*peak*_ and *V*_*AP*_ as well as between *T*_*APHW*_ and *dV*/*dt*|_*min*_. We found strong negative correlations between *dV*/*dt*|_*max*_ and *T*_*APHW*_ as well as between *dV*/*dt*|_*max*_ and *dV*/*dt*|_*min*_. These strong and weak correlations are along expected lines from the physiology of granule cells from intermediate sections (Mishra & Narayanan, 2020a). Together, the weak correlations spanning a majority of the measurement pairs indicated that the different measurements were quantifying distinct physiological characteristics of the granule cells across all three sections.

**Figure 5.**
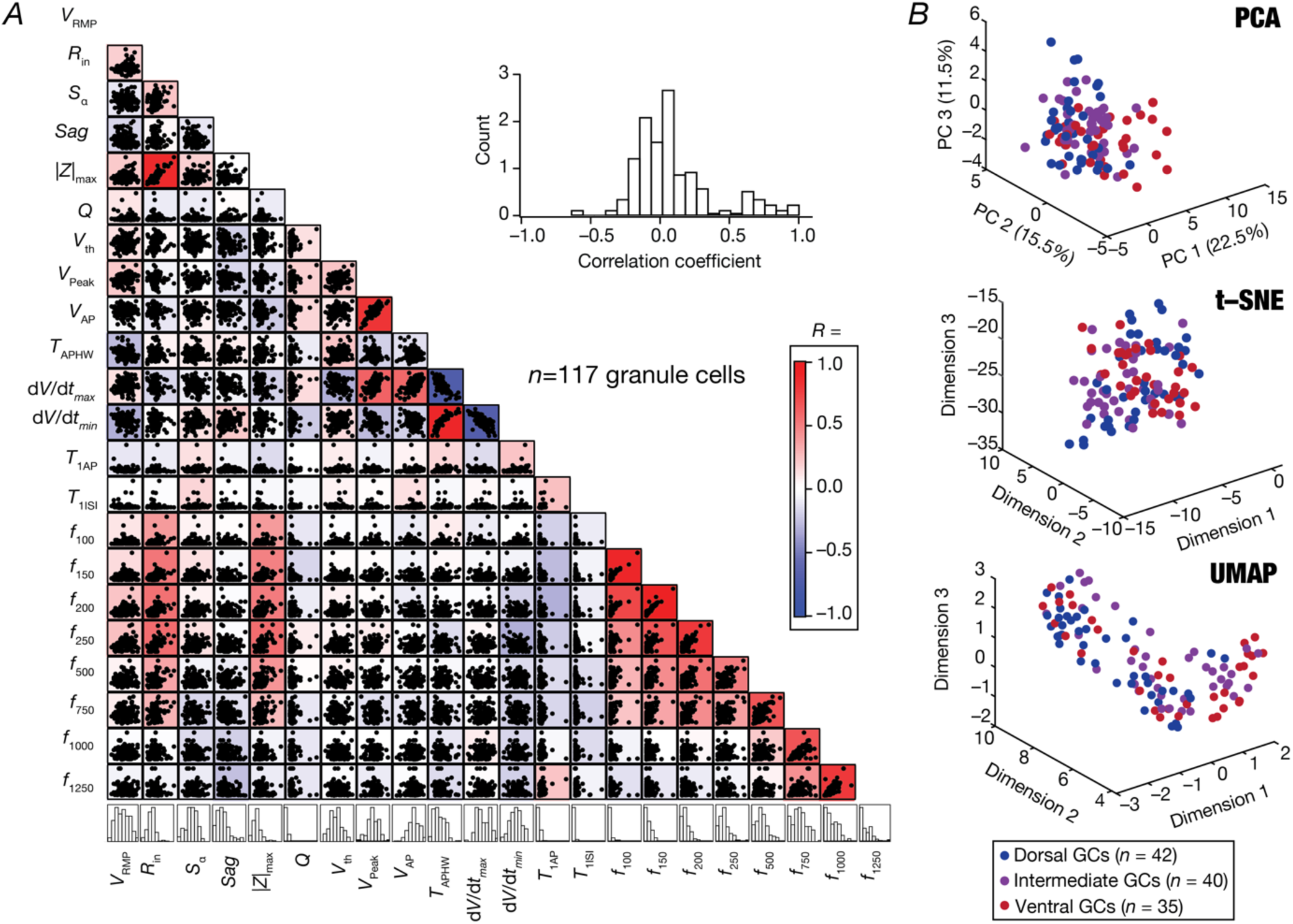
Correlation and dimensionality reduction analyses on the measurement space of granule cells across the hippocampal dorsoventral axis. *A,* Pairwise scatterplot matrices of 22 subthreshold and suprathreshold measurements of granule cells recorded from the dorsal, intermediate, and ventral regions pooled together. The scatterplot matrices are overlaid on the corresponding color-coded correlation matrices. The histograms of all the respective measurements are depicted at the bottom of the matrix. The inset represents the histogram of the correlation coefficients across all unique pairs of measurements. *B, Top*, Principal component analysis (PCA) on the 22-dimensional electrophysiological measurements of GCs across the dorsoventral axis of dentate gyrus. The percentage variance explained by each principal component are shown within parentheses in each axis label. *Middle*, Outcome of *t*-distributed stochastic neighbor embedding, *t*-SNE on the 22-dimensional measurement space. *Bottom*, uniform manifold approximation and projection, UMAP analyses on the 22-dimensional measurement space.

We asked whether DG granule-cell measurements form distinct dorsoventral groups and whether the measurements lie in a low-dimensional subspace. Specifically, were there distinct clusters for the dorsal (*n* = 42), intermediate (*n* = 40), and ventral (*n* = 35) DG GCs in the measurement space for the 117 neurons considered? Were there strong co-dependencies across the measurements, that are not observable as pairwise correlations but manifest as alignments in a low-dimensional subspace? How heterogeneous were the measurements in reduced dimensional spaces? Was the degree of heterogeneity dependent on the dorsoventral location of granule cells? To address these questions, we performed linear and non-linear dimensionality reduction analyses on the 22-dimensional measurement space. We did not consider resonance frequency (*f*_*R*_), total inductive phase (Φ_L_), and firing rate at 50 pA (*f*_KL_) because *f*_*R*_ was close to 1 and Φ_L_ and *f*_KL_ were close to zero across all neurons.

We applied principal component analysis (PCA), a linear dimensionality reduction analysis technique, on the 22-dimensional measurements from the three categories of GCs. We visualized the projections of the data onto the three principal dimensions, with labels identifying the dorsoventral localization of the individual neurons (Fig. 5*B*, *top*). We noted that the first three dimensions explained barely 50% of the variance in the data. These observations point to the absence of a low-dimensional subspace within the measurement space, consistent with our observation on the lack of strong correlations across a majority of measurement pairs (Fig. 5*A*). The high-dimensionality of the underlying data was further confirmed by the dimensionality of the space, computed as 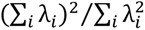 from the eigen values λ_*i*_, *i* = 1, …,22 (Abbott *et al*., 2011; Mazzucato *et al*., 2016; Litwin-Kumar *et al*., 2017), was 8.92. Second, although there were visible subregions containing predominantly dorsal or ventral neurons, strong overlaps were observed between the three different groups in the reduced dimensional space (Fig. 5*B*, *top*). These observations pointed to a lack of independent clusters in the measurement space specifically corresponding to the dorsal, intermediate, and ventral GCs. These observations are consistent with the widespread heterogeneities in each measurement across the three sections (Fig. 2, Fig. 4, Table 1) and the less number of measurements that were significantly different across sections (Fig. 2, Fig. 4, Table 2). We also performed nonlinear dimensionality reduction techniques *t*-SNE (Fig. 5*B*, *middle*) and UMAP (Fig. 5*B*, *bottom*) on the 22-dimensional measurement space associated with the 117 neurons. The outcomes were consistent with our observations from PCA, together confirming (i) the lack of strong cross-dependencies across the different measurements; and (ii) the absence of independent clusters for each of the three sections (Fig. 5*B*).

Together, pairwise correlation analyses and dimensionality reduction analyses showed that the measurement space lacked strong dependencies across the different physiological measurements. In addition, there were no independent clusters for each of the dorsal, ventral, and intermediate sections in the reduced dimensional spaces. The distribution of the data points and the variability measures (SEM, SD, IQR, and CV) associated with each measurement (Fig. 2, Fig. 4, Table 1, Tables 4–9) together emphasize the need to explicitly account for the widespread heterogeneities in physiological properties across the dorsoventral axis.

**Table 4:**
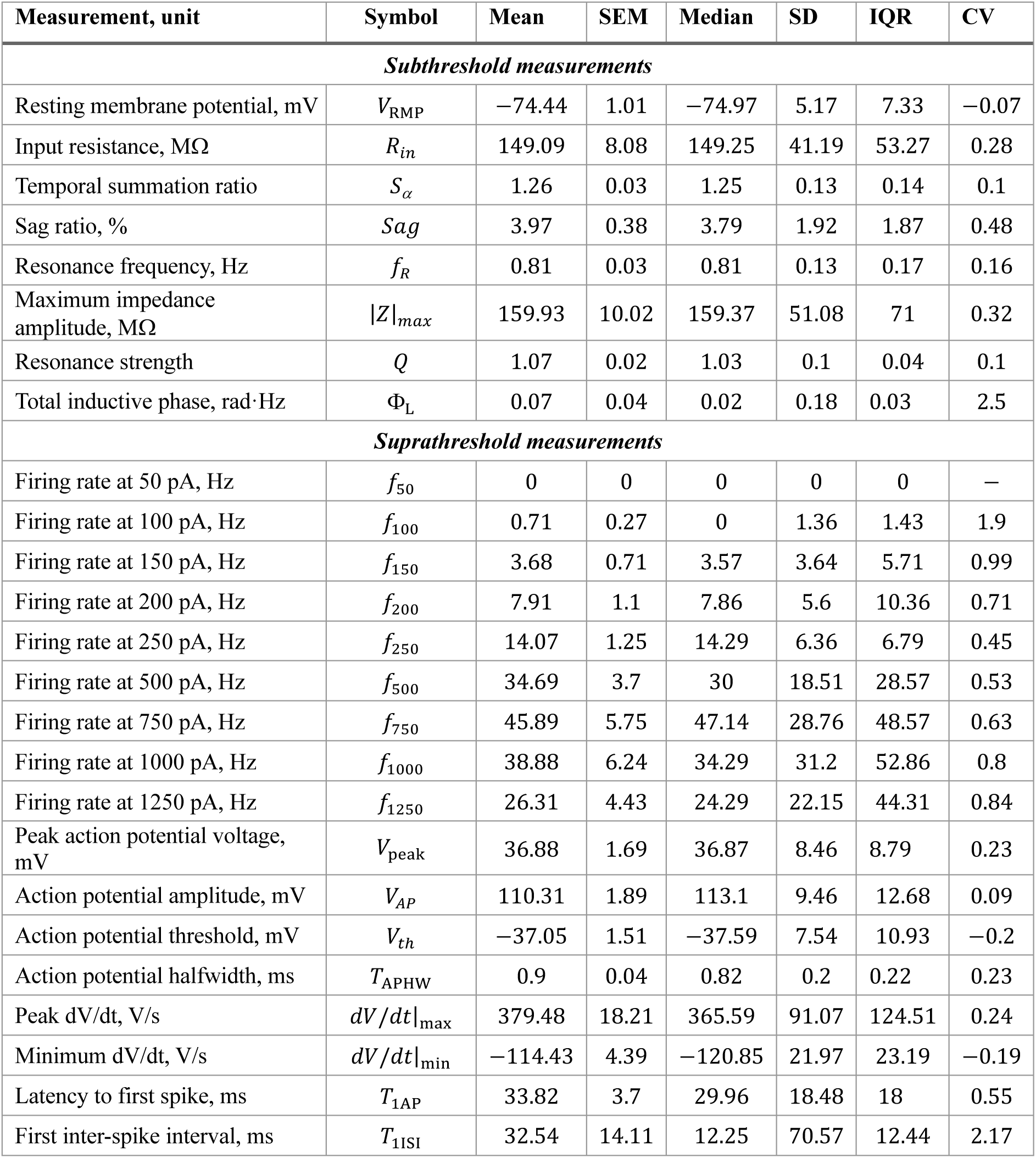
Summary statistics for subthreshold and suprathreshold measurements recorded from neurons in dorsal infrapyramidal GCs (*n* = 27) when respective current protocols were injected with cell resting at *V*_RMP_ (Fig. 6*B–G*). Measurements are mean, SEM, median number of cells and the degree of variability in each measurement is reported as standard deviation, interquartile range, and coefficient of variation, in that order.

| Measurement, unit | Symbol | Mean | SEM | Median | SD | IQR | CV |
| --- | --- | --- | --- | --- | --- | --- | --- |
| *Subthreshold measurements* |  |  |  |  |  |  |  |
| Resting membrane potential, mV | $V_{\text{RMP}}$ | −74.44 | 1.01 | −74.97 | 5.17 | 7.33 | −0.07 |
| Input resistance, MΩ | $R_{\text{in}}$ | 149.09 | 8.08 | 149.25 | 41.19 | 53.27 | 0.28 |
| Temporal summation ratio | $S_{\alpha}$ | 1.26 | 0.03 | 1.25 | 0.13 | 0.14 | 0.1 |
| Sag ratio, % | $S_{\text{ag}}$ | 3.97 | 0.38 | 3.79 | 1.92 | 1.87 | 0.48 |
| Resonance frequency, Hz | $f_{\text{R}}$ | 0.81 | 0.03 | 0.81 | 0.13 | 0.17 | 0.16 |
| Maximum impedance amplitude, MΩ | $\vert Z\vert _{\text{max}}$ | 159.93 | 10.02 | 159.37 | 51.08 | 71 | 0.32 |
| Resonance strength | $Q$ | 1.07 | 0.02 | 1.03 | 0.1 | 0.04 | 0.1 |
| Total inductive phase, rad·Hz | $\Phi_{\text{L}}$ | 0.07 | 0.04 | 0.02 | 0.18 | 0.03 | 2.5 |
| *Suprathreshold measurements* |  |  |  |  |  |  |  |
| Firing rate at 50 pA, Hz | $f_{50}$ | 0 | 0 | 0 | 0 | 0 | — |
| Firing rate at 100 pA, Hz | $f_{100}$ | 0.71 | 0.27 | 0 | 1.36 | 1.43 | 1.9 |
| Firing rate at 150 pA, Hz | $f_{150}$ | 3.68 | 0.71 | 3.57 | 3.64 | 5.71 | 0.99 |
| Firing rate at 200 pA, Hz | $f_{200}$ | 7.91 | 1.1 | 7.86 | 5.6 | 10.36 | 0.71 |
| Firing rate at 250 pA, Hz | $f_{250}$ | 14.07 | 1.25 | 14.29 | 6.36 | 6.79 | 0.45 |
| Firing rate at 500 pA, Hz | $f_{500}$ | 34.69 | 3.7 | 30 | 18.51 | 28.57 | 0.53 |
| Firing rate at 750 pA, Hz | $f_{750}$ | 45.89 | 5.75 | 47.14 | 28.76 | 48.57 | 0.63 |
| Firing rate at 1000 pA, Hz | $f_{1000}$ | 38.88 | 6.24 | 34.29 | 31.2 | 52.86 | 0.8 |
| Firing rate at 1250 pA, Hz | $f_{1250}$ | 26.31 | 4.43 | 24.29 | 22.15 | 44.31 | 0.84 |
| Peak action potential voltage, mV | $V_{\text{peak}}$ | 36.88 | 1.69 | 36.87 | 8.46 | 8.79 | 0.23 |
| Action potential amplitude, mV | $V_{\text{AP}}$ | 110.31 | 1.89 | 113.1 | 9.46 | 12.68 | 0.09 |
| Action potential threshold, mV | $V_{\text{th}}$ | −37.05 | 1.51 | −37.59 | 7.54 | 10.93 | −0.2 |
| Action potential halfwidth, ms | $T_{\text{APHW}}$ | 0.9 | 0.04 | 0.82 | 0.2 | 0.22 | 0.23 |
| Peak dV/dt, V/s | $dV/dt\vert _{\text{max}}$ | 379.48 | 18.21 | 365.59 | 91.07 | 124.51 | 0.24 |
| Minimum dV/dt, V/s | $dV/dt\vert _{\text{min}}$ | −114.43 | 4.39 | −120.85 | 21.97 | 23.19 | −0.19 |
| Latency to first spike, ms | $T_{1\text{AP}}$ | 33.82 | 3.7 | 29.96 | 18.48 | 18 | 0.55 |
| First inter-spike interval, ms | $T_{1\text{ISI}}$ | 32.54 | 14.11 | 12.25 | 70.57 | 12.44 | 2.17 |

**Table 5:** Summary statistics for subthreshold and suprathreshold measurements recorded from neurons in dorsal suprapyramidal GCs (*n* = 24) when respective current protocols were injected with cell resting at *V*_RMP_ (Fig. 6*B–G*). Measurements are mean, SEM, median number of cells and the degree of variability in each measurement is reported as standard deviation, interquartile range, and coefficient of variation.

| Measurement, unit | Symbol | Mean | SEM | Median | SD | IQR | CV |
| --- | --- | --- | --- | --- | --- | --- | --- |
| *Subthreshold measurements* |  |  |  |  |  |  |  |
| Resting membrane potential, mV | $V_{RMP}$ | −70.91 | 0.93 | −70.6 | 4.48 | 6.51 | −0.06 |
| Input resistance, MΩ | $R_{in}$ | 118.14 | 11.25 | 130.15 | 53.97 | 65.13 | 0.46 |
| Temporal summation ratio | $S_{\alpha}$ | 1.28 | 0.02 | 1.31 | 0.11 | 0.1 | 0.09 |
| Sag ratio, % | $Sag$ | 3.56 | 0.5 | 3.77 | 2.31 | 4.14 | 0.65 |
| Resonance frequency, Hz | $f_R$ | 0.75 | 0.04 | 0.83 | 0.19 | 0.3 | 0.26 |
| Maximum impedance amplitude, MΩ | $\vert Z\vert _{max}$ | 152.52 | 11.91 | 153.32 | 54.58 | 66.23 | 0.36 |
| Resonance strength | $Q$ | 1.04 | 0.01 | 1.02 | 0.04 | 0.03 | 0.03 |
| Total inductive phase, rad·Hz | $\Phi_L$ | 0.11 | 0.08 | 0 | 0.37 | 0.04 | 3.38 |
| *Suprathreshold measurements* |  |  |  |  |  |  |  |
| Firing rate at 50 pA, Hz | $f_{50}$ | 0 | 0 | 0 | 0 | 0 | — |
| Firing rate at 100 pA, Hz | $f_{100}$ | 0.68 | 0.22 | 0 | 1.04 | 1.43 | 1.53 |
| Firing rate at 150 pA, Hz | $f_{150}$ | 3.48 | 0.66 | 4.29 | 3.16 | 5 | 0.91 |
| Firing rate at 200 pA, Hz | $f_{200}$ | 7.08 | 1.22 | 7.14 | 5.85 | 7.86 | 0.83 |
| Firing rate at 250 pA, Hz | $f_{250}$ | 10.68 | 1.57 | 10 | 7.53 | 9.29 | 0.71 |
| Firing rate at 500 pA, Hz | $f_{500}$ | 25.08 | 4.24 | 26.43 | 17.99 | 24.64 | 0.72 |
| Firing rate at 750 pA, Hz | $f_{750}$ | 33.87 | 4.73 | 38.57 | 19.51 | 24.29 | 0.58 |
| Firing rate at 1000 pA, Hz | $f_{1000}$ | 29.65 | 5.01 | 31.43 | 20.68 | 22.86 | 0.7 |
| Firing rate at 1250 pA, Hz | $f_{1250}$ | 19.75 | 4.04 | 17.14 | 16.65 | 18.57 | 0.84 |
| Peak action potential voltage, mV | $V_{peak}$ | 44.75 | 1.72 | 45.7 | 7.71 | 8.09 | 0.17 |
| Action potential amplitude, mV | $V_{AP}$ | 114.91 | 2.04 | 115.37 | 9.11 | 13.04 | 0.08 |
| Action potential threshold, mV | $V_{th}$ | −38.2 | 1.65 | −37.65 | 7.38 | 10.16 | −0.19 |
| Action potential halfwidth, ms | $T_{APHW}$ | 0.83 | 0.07 | 0.73 | 0.3 | 0.24 | 0.36 |
| Peak dV/dt, V/s | $dV/dt\vert _{max}$ | 447.77 | 18.44 | 465.99 | 82.48 | 50.66 | 0.18 |
| Minimum dV/dt, V/s | $dV/dt\vert _{min}$ | −129.76 | 7.13 | −137.94 | 31.9 | 35.09 | −0.25 |
| Latency to first spike, ms | $T_{1AP}$ | 23.16 | 3.86 | 24.09 | 17.26 | 27.34 | 0.75 |
| First inter-spike interval, ms | $T_{1ISI}$ | 11.88 | 1.79 | 6.86 | 7.99 | 14.78 | 0.67 |

**Table 6:** Summary statistics for subthreshold and suprathreshold measurements recorded from neurons in intermediate infrapyramidal GCs (*n* = 21) when respective current protocols were injected with cell resting at *V*_RMP_ (Fig. 7*B–G*). Measurements are mean, SEM, median number of cells and the degree of variability in each measurement is reported as standard deviation, interquartile range, and coefficient of variation.

| Measurement, unit | Symbol | Mean | SEM | Median | SD | IQR | CV |
| --- | --- | --- | --- | --- | --- | --- | --- |
| *Subthreshold measurements* |  |  |  |  |  |  |  |
| Resting membrane potential, mV | $V_{RMP}$ | −71.02 | 1.66 | −71.72 | 7.6 | 12.68 | −0.11 |
| Input resistance, MΩ | $R_{in}$ | 167.98 | 16.7 | 151.11 | 76.51 | 59.63 | 0.46 |
| Temporal summation ratio | $S_{\alpha}$ | 1.32 | 0.03 | 1.33 | 0.16 | 0.14 | 0.12 |
| Sag ratio, % | $Sag$ | 3.68 | 0.6 | 2.62 | 2.77 | 2.76 | 0.75 |
| Resonance frequency, Hz | $f_R$ | 0.82 | 0.04 | 0.8 | 0.2 | 0.25 | 0.25 |
| Maximum impedance amplitude, MΩ | $\vert Z\vert _{max}$ | 185.99 | 19.57 | 170.82 | 89.7 | 72.05 | 0.48 |
| Resonance strength | $Q$ | 1.09 | 0.03 | 1.04 | 0.15 | 0.05 | 0.14 |
| Total inductive phase, rad·Hz | $\Phi_L$ | 0.04 | 0.01 | 0.01 | 0.06 | 0.06 | 1.74 |
| *Suprathreshold measurements* |  |  |  |  |  |  |  |
| Firing rate at 50 pA, Hz | $f_{50}$ | 0.07 | 0.07 | 0 | 0.31 | 0 | 4.58 |
| Firing rate at 100 pA, Hz | $f_{100}$ | 0.48 | 0.35 | 0 | 1.59 | 0 | 3.33 |
| Firing rate at 150 pA, Hz | $f_{150}$ | 5.17 | 1.1 | 5.71 | 5.06 | 5.71 | 0.98 |
| Firing rate at 200 pA, Hz | $f_{200}$ | 12.38 | 2.32 | 11.43 | 10.62 | 8.57 | 0.86 |
| Firing rate at 250 pA, Hz | $f_{250}$ | 20.2 | 3.25 | 17.14 | 14.91 | 10 | 0.74 |
| Firing rate at 500 pA, Hz | $f_{500}$ | 48.2 | 6.59 | 42.86 | 28.74 | 20.71 | 0.6 |
| Firing rate at 750 pA, Hz | $f_{750}$ | 59.01 | 8.56 | 51.43 | 36.31 | 37.86 | 0.62 |
| Firing rate at 1000 pA, Hz | $f_{1000}$ | 44.68 | 9.42 | 30.71 | 39.95 | 48.21 | 0.89 |
| Firing rate at 1250 pA, Hz | $f_{1250}$ | 30.08 | 5.86 | 18.57 | 24.87 | 42.5 | 0.83 |
| Peak action potential voltage, mV | $V_{peak}$ | 42.72 | 2.63 | 45.93 | 12.05 | 14.71 | 0.28 |
| Action potential amplitude, mV | $V_{AP}$ | 113.72 | 2.07 | 112.52 | 9.51 | 9.5 | 0.08 |
| Action potential threshold, mV | $V_{th}$ | −33.32 | 1.43 | −34.25 | 6.56 | 6.01 | −0.2 |
| Action potential halfwidth, ms | $T_{APHW}$ | 0.95 | 0.04 | 0.95 | 0.17 | 0.24 | 0.18 |
| Peak dV/dt, V/s | $dV/dt\vert _{max}$ | 363.7 | 18.01 | 349.72 | 82.55 | 100.71 | 0.23 |
| Minimum dV/dt, V/s | $dV/dt\vert _{min}$ | −112.94 | 5.94 | −108.03 | 27.22 | 24.41 | −0.24 |
| Latency to first spike, ms | $T_{1AP}$ | 48.87 | 8.64 | 43.7 | 39.59 | 24.65 | 0.81 |
| First inter-spike interval, ms | $T_{1ISI}$ | 32.55 | 9.79 | 17 | 44.86 | 26.92 | 1.38 |

**Table 7:** Summary statistics for subthreshold and suprathreshold measurements recorded from neurons in intermediate suprapyramidal GCs (*n* = 21) when respective current protocols were injected with cell resting at *V*_RMP_ (Fig. 7*B–G*). Measurements are mean, SEM, median number of cells and the degree of variability in each measurement is reported as standard deviation, interquartile range, and coefficient of variation.

| Measurement, unit | Symbol | Mean | SEM | Median | SD | IQR | CV |
| --- | --- | --- | --- | --- | --- | --- | --- |
| *Subthreshold measurements* |  |  |  |  |  |  |  |
| Resting membrane potential, mV | $V_{RMP}$ | −72.96 | 1.29 | −73.73 | 5.93 | 9.64 | −0.08 |
| Input resistance, MΩ | $R_{in}$ | 174.36 | 15.34 | 182.37 | 68.62 | 77.96 | 0.39 |
| Temporal summation ratio | $S_{\alpha}$ | 1.23 | 0.02 | 1.22 | 0.11 | 0.14 | 0.09 |
| Sag ratio, % | $Sag$ | 3.05 | 0.41 | 2.53 | 1.89 | 2.32 | 0.62 |
| Resonance frequency, Hz | $f_R$ | 0.77 | 0.04 | 0.74 | 0.18 | 0.27 | 0.23 |
| Maximum impedance amplitude, MΩ | $\vert Z\vert _{max}$ | 192.2 | 15.4 | 191.1 | 70.7 | 88.7 | 0.37 |
| Resonance strength | $Q$ | 1.05 | 0.01 | 1.03 | 0.06 | 0.03 | 0.06 |
| Total inductive phase, rad·Hz | $\Phi_L$ | 0.03 | 0.01 | 0.01 | 0.07 | 0.04 | 1.98 |
| *Suprathreshold measurements* |  |  |  |  |  |  |  |
| Firing rate at 50 pA, Hz | $f_{50}$ | 0.21 | 0.21 | 0 | 0.96 | 0 | 4.47 |
| Firing rate at 100 pA, Hz | $f_{100}$ | 2.36 | 0.8 | 0 | 3.6 | 3.21 | 1.53 |
| Firing rate at 150 pA, Hz | $f_{150}$ | 7.71 | 1.43 | 7.86 | 6.4 | 8.57 | 0.83 |
| Firing rate at 200 pA, Hz | $f_{200}$ | 13.5 | 2.04 | 14.29 | 9.11 | 12.14 | 0.67 |
| Firing rate at 250 pA, Hz | $f_{250}$ | 18.23 | 2.69 | 20 | 12.33 | 12.86 | 0.68 |
| Firing rate at 500 pA, Hz | $f_{500}$ | 53.16 | 5.99 | 55.71 | 27.43 | 38.57 | 0.52 |
| Firing rate at 750 pA, Hz | $f_{750}$ | 72.17 | 5.56 | 68.57 | 25.5 | 30 | 0.35 |
| Firing rate at 1000 pA, Hz | $f_{1000}$ | 72.04 | 6.06 | 74.29 | 27.78 | 35.71 | 0.39 |
| Firing rate at 1250 pA, Hz | $f_{1250}$ | 51.84 | 7.62 | 52.86 | 34.91 | 30 | 0.67 |
| Peak action potential voltage, mV | $V_{peak}$ | 42.71 | 1.71 | 41.6 | 7.47 | 11.38 | 0.17 |
| Action potential amplitude, mV | $V_{AP}$ | 115.78 | 1.28 | 116.55 | 5.59 | 4.35 | 0.05 |
| Action potential threshold, mV | $V_{th}$ | −35.67 | 1.75 | −35.5 | 7.63 | 8.59 | −0.21 |
| Action potential halfwidth, ms | $T_{APHW}$ | 1.08 | 0.06 | 1.14 | 0.24 | 0.37 | 0.22 |
| Peak dV/dt, V/s | $dV/dt\vert _{max}$ | 350.72 | 24.76 | 313.7 | 107.92 | 125.73 | 0.31 |
| Minimum dV/dt, V/s | $dV/dt\vert _{min}$ | −107.39 | 5.36 | −100.7 | 23.37 | 16.78 | −0.22 |
| Latency to first spike, ms | $T_{1AP}$ | 30.43 | 4.8 | 28.58 | 20.91 | 24.48 | 0.69 |
| First inter-spike interval, ms | $T_{1ISI}$ | 9.49 | 1.84 | 7.34 | 7.58 | 2.92 | 0.8 |

**Table 8:** Summary statistics for subthreshold and suprathreshold measurements recorded from neurons in ventral infrapyramidal GCs (*n* = 23) when respective current protocols were injected with cell resting at *V*_RMP_(Fig. 8*B–G*). Measurements are mean, SEM, median number of cells and the degree of variability in each measurement is reported as standard deviation, interquartile range, and coefficient of variation.

| Measurement, unit | Symbol | Mean | SEM | Median | SD | IQR | CV |
| --- | --- | --- | --- | --- | --- | --- | --- |
| *Subthreshold measurements* |  |  |  |  |  |  |  |
| Resting membrane potential, mV | $V_{RMP}$ | −71.4 | 0.91 | −71.6 | 4.40 | 6.31 | −0.06 |
| Input resistance, MΩ | $R_{in}$ | 215.16 | 19.20 | 174.7 | 92.11 | 92.10 | 0.42 |
| Temporal summation ratio | $S_{\alpha}$ | 1.30 | 0.02 | 1.31 | 0.12 | 0.11 | 0.09 |
| Sag ratio, % | $Sag$ | 4.12 | 0.47 | 3.52 | 2.29 | 2.60 | 0.55 |
| Resonance frequency, Hz | $f_R$ | 0.73 | 0.04 | 0.74 | 0.20 | 0.24 | 0.27 |
| Maximum impedance amplitude, MΩ | $\vert Z\vert _{max}$ | 244.6 | 26.85 | 185.92 | 128.79 | 147.98 | 0.52 |
| Resonance strength | $Q$ | 1.03 | 0.005 | 1.03 | 0.02 | 0.03 | 0.025 |
| Total inductive phase, rad·Hz | $\Phi_L$ | 0.03 | 0.018 | 0.004 | 0.09 | 0.02 | 2.57 |
| *Suprathreshold measurements* |  |  |  |  |  |  |  |
| Firing rate at 50 pA, Hz | $f_{50}$ | 1.82 | 0.93 | 0 | 4.37 | 0 | 2.4 |
| Firing rate at 100 pA, Hz | $f_{100}$ | 6.32 | 2.09 | 2.86 | 9.81 | 8.57 | 1.55 |
| Firing rate at 150 pA, Hz | $f_{150}$ | 14.31 | 2.54 | 12.86 | 11.89 | 9.64 | 0.83 |
| Firing rate at 200 pA, Hz | $f_{200}$ | 20.43 | 2.95 | 20.71 | 13.85 | 11.43 | 0.68 |
| Firing rate at 250 pA, Hz | $f_{250}$ | 26.9 | 2.94 | 27.86 | 13.81 | 11.43 | 0.51 |
| Firing rate at 500 pA, Hz | $f_{500}$ | 55.25 | 6.38 | 45 | 27.05 | 44.64 | 0.49 |
| Firing rate at 750 pA, Hz | $f_{750}$ | 64.83 | 7.48 | 62.14 | 31.74 | 34.64 | 0.49 |
| Firing rate at 1000 pA, Hz | $f_{1000}$ | 42.51 | 6.3 | 45 | 26.75 | 39.8 | 0.63 |
| Firing rate at 1250 pA, Hz | $f_{1250}$ | 21.85 | 4.05 | 15.71 | 17.2 | 27.19 | 0.79 |
| Peak action potential voltage, mV | $V_{peak}$ | 44.73 | 2.34 | 44.63 | 10.98 | 12.36 | 0.25 |
| Action potential amplitude, mV | $V_{AP}$ | 115.01 | 2.15 | 116.71 | 10.07 | 12.16 | 0.09 |
| Action potential threshold, mV | $V_{th}$ | −34.81 | 1.74 | −35.8 | 8.15 | 14.83 | −0.23 |
| Action potential halfwidth, ms | $T_{APHW}$ | 1.03 | 0.05 | 1.03 | 0.25 | 0.28 | 0.25 |
| Peak dV/dt, V/s | $dV/dt\vert _{max}$ | 381.2 | 25.93 | 383.3 | 121.6 | 185.4 | 0.32 |
| Minimum dV/dt, V/s | $dV/dt\vert _{min}$ | −108.9 | 6.1 | −106.5 | 28.61 | 33.7 | −0.26 |
| Latency to first spike, ms | $T_{1AP}$ | 28.22 | 5.2 | 23.8 | 24.38 | 19.9 | 0.86 |
| First inter-spike interval, ms | $T_{1ISI}$ | 10.84 | 2.8 | 6.07 | 12.83 | 2.9 | 1.18 |

**Table 9:** Summary statistics for subthreshold and suprathreshold measurements recorded from neurons in ventral suprapyramidal GCs (*n* = 20) when respective current protocols were injected with cell resting at *V*_RMP_ (Fig. 8*B–G*). Measurements are mean, SEM, median number of cells and the degree of variability in each measurement is reported as standard deviation, interquartile range, and coefficient of variation.

| Measurement, unit | Symbol | Mean | SEM | Median | SD | IQR | CV |
| --- | --- | --- | --- | --- | --- | --- | --- |
| *Subthreshold measurements* |  |  |  |  |  |  |  |
| Resting membrane potential, mV | $V_{RMP}$ | −72.02 | 0.67 | −73.2 | 3.01 | 5.2 | −0.04 |
| Input resistance, MΩ | $R_{in}$ | 173.53 | 13.13 | 165.04 | 58.73 | 65.77 | 0.34 |
| Temporal summation ratio | $S_{\alpha}$ | 1.31 | 0.04 | 1.27 | 0.16 | 0.22 | 0.12 |
| Sag ratio, % | $Sag$ | 2.99 | 0.43 | 2.86 | 1.92 | 1.49 | 0.64 |
| Resonance frequency, Hz | $f_R$ | 0.87 | 0.05 | 0.8 | 0.23 | 0.2 | 0.27 |
| Maximum impedance amplitude, MΩ | $\vert Z\vert _{max}$ | 209.58 | 17.71 | 195.93 | 79.2 | 120.16 | 0.38 |
| Resonance strength | $Q$ | 1.05 | 0.01 | 1.05 | 0.04 | 0.06 | 0.04 |
| Total inductive phase, rad·Hz | $\Phi_L$ | 0.04 | 0.01 | 0 | 0.06 | 0.03 | 1.81 |
| *Suprathreshold measurements* |  |  |  |  |  |  |  |
| Firing rate at 50 pA, Hz | $f_{50}$ | 0.23 | 0.23 | 0 | 0.98 | 0 | 4.36 |
| Firing rate at 100 pA, Hz | $f_{100}$ | 1.28 | 0.79 | 0 | 3.43 | 0 | 2.68 |
| Firing rate at 150 pA, Hz | $f_{150}$ | 4.96 | 1.29 | 2.86 | 5.6 | 7.14 | 1.13 |
| Firing rate at 200 pA, Hz | $f_{200}$ | 9.7 | 1.69 | 8.57 | 7.36 | 8.57 | 0.76 |
| Firing rate at 250 pA, Hz | $f_{250}$ | 14.79 | 1.85 | 13.57 | 8.27 | 7.5 | 0.56 |
| Firing rate at 500 pA, Hz | $f_{500}$ | 32.3 | 3.99 | 31.43 | 16.93 | 22.5 | 0.52 |
| Firing rate at 750 pA, Hz | $f_{750}$ | 44.71 | 6.27 | 41.43 | 25.86 | 35.71 | 0.58 |
| Firing rate at 1000 pA, Hz | $f_{1000}$ | 34.29 | 5.5 | 32.86 | 22.67 | 22.86 | 0.66 |
| Firing rate at 1250 pA, Hz | $f_{1250}$ | 26.52 | 5.81 | 15.71 | 23.23 | 31.07 | 0.88 |
| Peak action potential voltage, mV | $V_{peak}$ | 39.65 | 2.19 | 42.48 | 9.53 | 8.96 | 0.24 |
| Action potential amplitude, mV | $V_{AP}$ | 111.27 | 2.48 | 112.57 | 10.82 | 13.95 | 0.1 |
| Action potential threshold, mV | $V_{th}$ | −31.09 | 1.88 | −31.47 | 8.22 | 12.53 | −0.26 |
| Action potential halfwidth, ms | $T_{APHW}$ | 1.07 | 0.08 | 1.01 | 0.34 | 0.43 | 0.32 |
| Peak dV/dt, V/s | $dV/dt\vert _{max}$ | 332.73 | 31.76 | 312.49 | 138.44 | 164.18 | 0.42 |
| Minimum dV/dt, V/s | $dV/dt\vert _{min}$ | −99.58 | 6.57 | −98.87 | 28.64 | 45.47 | −0.29 |
| Latency to first spike, ms | $T_{1AP}$ | 59.84 | 20.06 | 38.14 | 87.43 | 22.79 | 1.46 |
| First inter-spike interval, ms | $T_{1ISI}$ | 32.64 | 12.14 | 11.27 | 51.51 | 19.36 | 1.58 |

### Ventral granule cells in the infrapyramidal blade manifested higher firing rates compared to those in the suprapyramidal blade

Within each section along the dorsoventral axis, the dentate gyrus is anatomically further divided into two blades (Amaral *et al*., 2007). The granule cell layer of the DG closer to CA1 is referred to as the suprapyramidal blade (SPB) and the one farther away from CA1 was referred to as the infrapyramidal blade (IPB) (Fig. 6*A*). Are there blade-specific differences in intrinsic properties of granule cells located within each of the three sections? To address this, we identified granule cells based on their blade localization and compared their sub- and Suprathreshold physiological properties between SPB and IPB of dorsal (Fig. 6; Tables 4–5), intermediate (Fig. 7; Tables 6–7), and ventral (Fig. 8; Tables 8–9) dentate gyrus.

**Figure 6:**
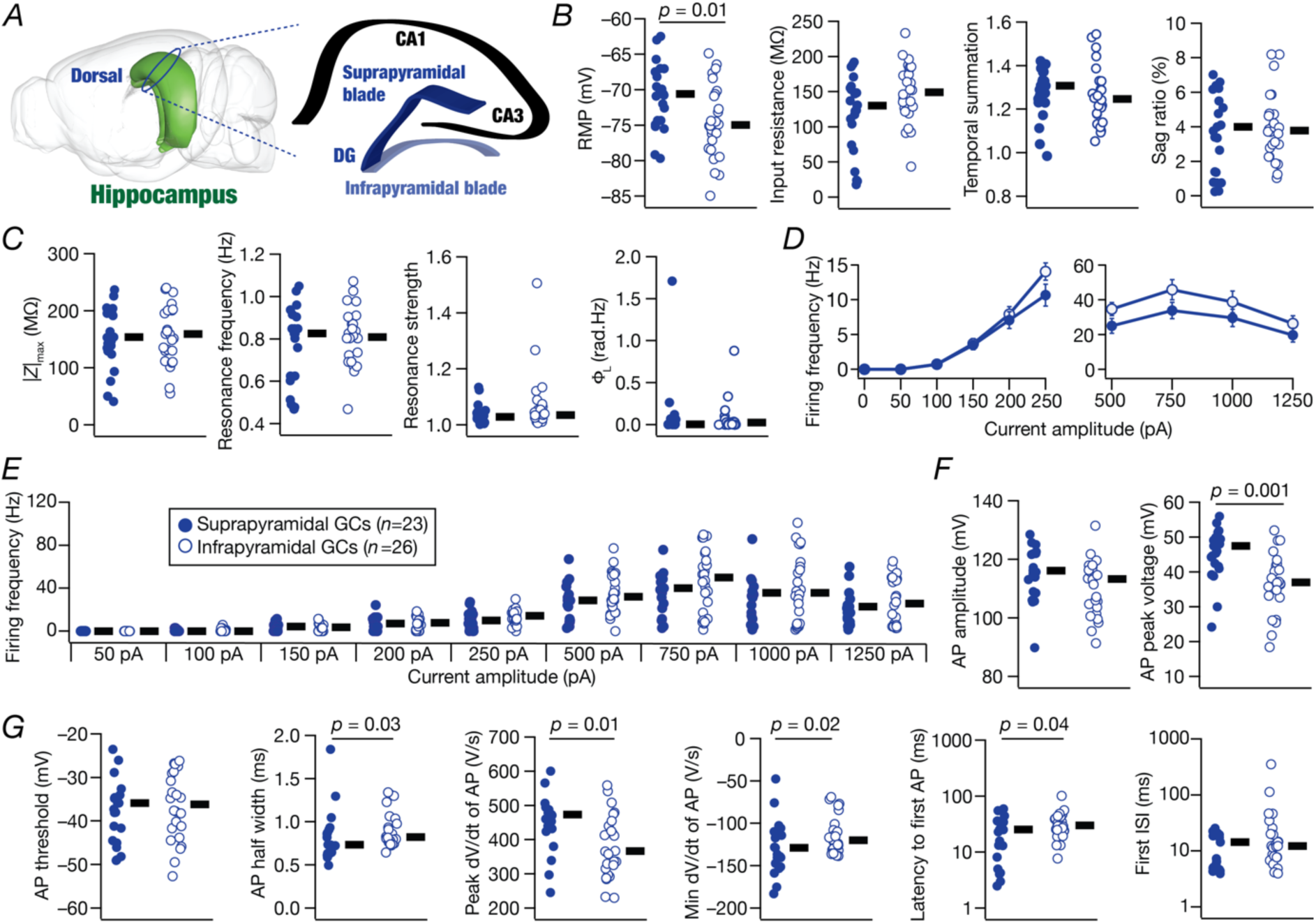
Heterogeneities in electrophysiological properties of granule cells across the blades of the dorsal dentate gyrus. *A*, Schematic of rodent brain showing the anatomical structure of a section of dorsal hippocampus (Image from Scalable Brain Atlas (Bakker *et al*., 2015)). *B–C,* Beeswarm plots comparing the subthreshold properties of dorsal granule cells from the infrapyramidal (open circles) and suprapyramidal (closed circles) blades. *D, f* − *I* curves of action potential (AP) firing rate *vs.* injected current amplitude, plotted for lower current injections 50 to 250 pA in steps of 50 pA (left) and for higher current injections from 500 pA to 1250 pA in steps of 250 pA (right). Comparisons are presented for dorsal granule cells from the suprapyramidal blade (closed circles, *n* = 23) *vs.* infrapyramidal blade (open circles, *n* = 26). *E,* Beeswarm plots of firing frequencies for all measured current injections for each measured cell, comparing dorsal suprapyramidal and infrapyramidal granule cells. *F–G*, Beeswarm plots comparing action potential properties of granule cells from both dorsal blades. *Statistical test: Wilcoxon rank sum test was performed between the groups, significant p values are mentioned on the top of figures. Additional summary statistics are provided in Tables 4–5. All statistical test outcomes associated with this figure are provided in Table 10*.

**Figure 7:**
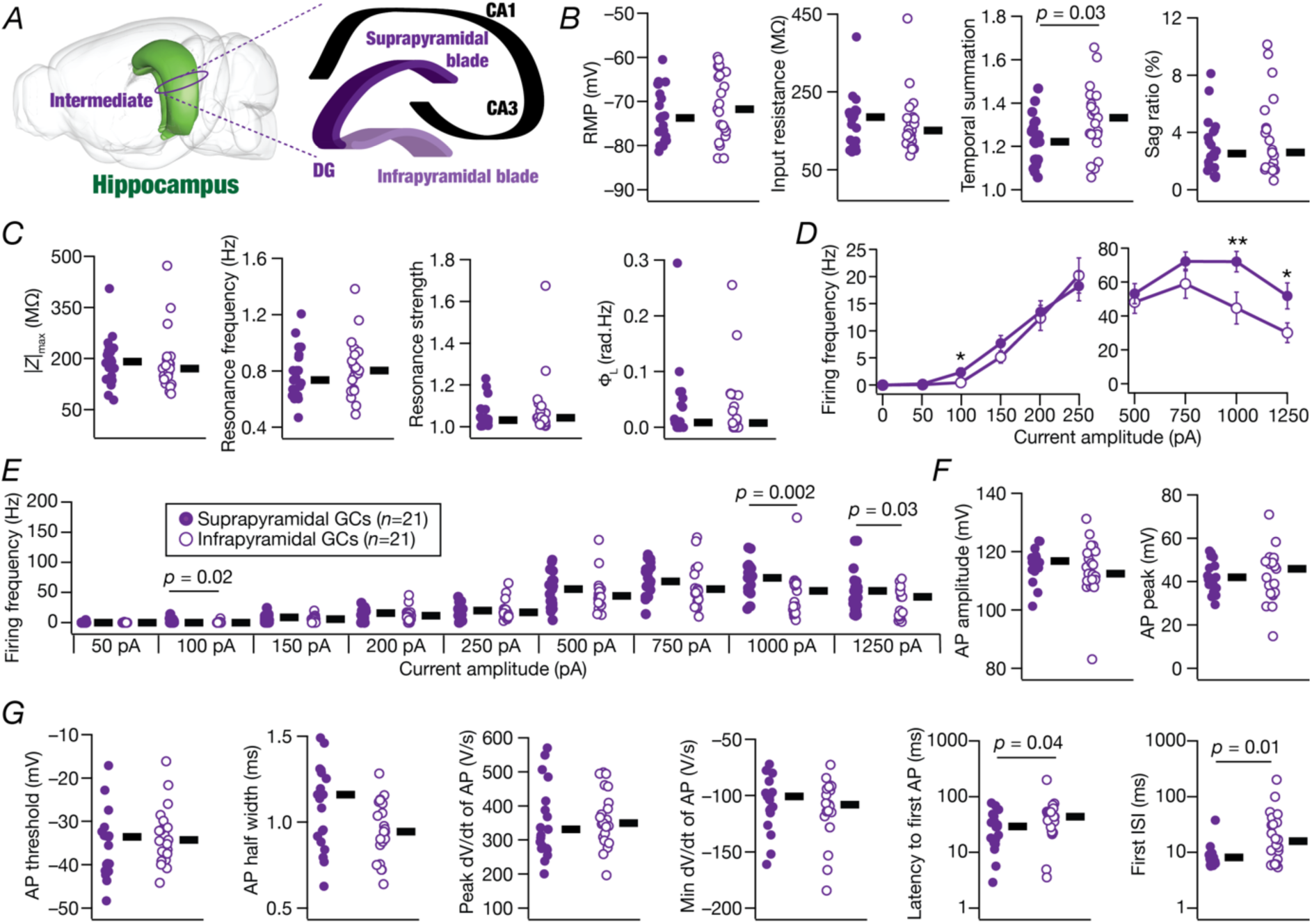
Heterogeneities in electrophysiological properties of granule cells across the blades of the intermediate dentate gyrus. *A*, Schematic of rodent brain showing the anatomical structure of a section of intermediate hippocampus (Image from Scalable Brain Atlas (Bakker *et al*., 2015)). *B–C,* Beeswarm plots comparing the subthreshold properties of intermediate granule cells from the infrapyramidal (open circles) and suprapyramidal (closed circles) blades. *D, f* − *I* curves of action potential (AP) firing rate *vs.* injected current amplitude, plotted for lower current injections 50 to 250 pA in steps of 50 pA (left) and for higher current injections from 500 pA to 1250 pA in steps of 250 pA (right). Comparisons are presented for intermediate granule cells from the suprapyramidal blade (closed circles, *n* = 21) *vs.* infrapyramidal blade (open circles, *n* = 21). *p values: * <0.05, ** < 0.005, *** < 0.0005*. *E,* Beeswarm plots of firing frequencies for all measured current injections for each measured cell, comparing intermediate suprapyramidal and infrapyramidal granule cells. *F– G*, Beeswarm plots comparing action potential properties of granule cells from both intermediate blades. *Statistical test: Wilcoxon rank sum test was performed between the groups, significant p values are mentioned on the top of figures. Additional summary statistics are provided in Tables 6–7. All statistical test outcomes associated with this figure are provided in Table 10*.

**Figure 8:**
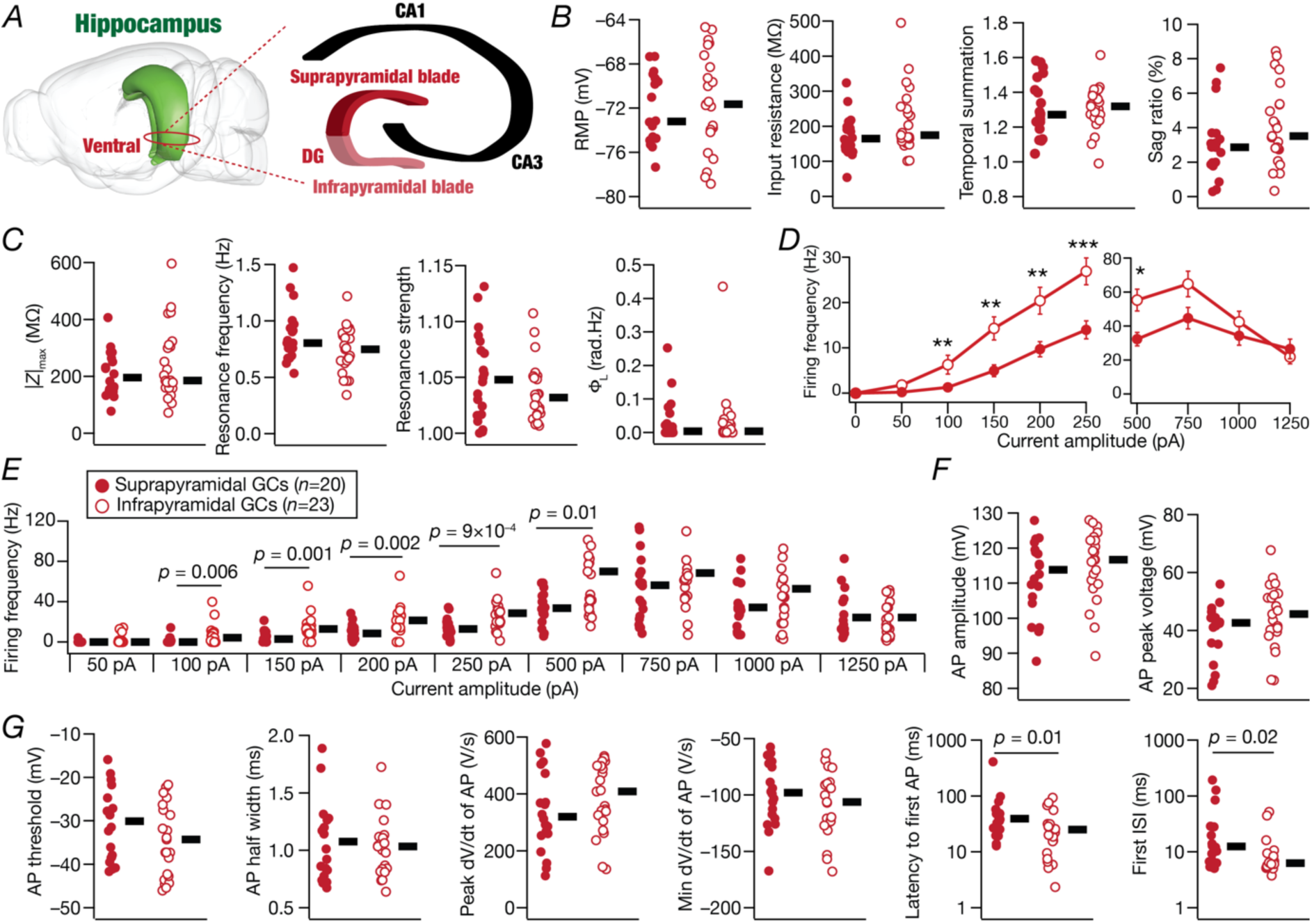
Heterogeneities in electrophysiological properties of granule cells across the blades of the ventral dentate gyrus. *A*, Schematic of rodent brain showing the anatomical structure of a section of ventral hippocampus (Image from Scalable Brain Atlas (Bakker *et al*., 2015)). *B–C,* Beeswarm plots comparing the subthreshold properties of ventral granule cells from the infrapyramidal (open circles) and suprapyramidal (closed circles) blades. *D, f* − *I* curves of action potential (AP) firing rate *vs.* injected current amplitude, plotted for lower current injections 50 to 250 pA in steps of 50 pA (left) and for higher current injections from 500 pA to 1250 pA in steps of 250 pA (right). Comparisons are presented for ventral granule cells from the suprapyramidal blade (closed circles, *n* = 20) *vs.* infrapyramidal blade (open circles, *n* = 23). *p values: * <0.05, ** < 0.005, *** < 0.0005*. *E,* Beeswarm plots of firing frequencies for all measured current injections for each measured cell, comparing ventral suprapyramidal and infrapyramidal granule cells. *F–G*, Beeswarm plots comparing action potential properties of granule cells from both ventral blades. *Statistical test: Wilcoxon rank sum test was performed between the groups, significant p values are mentioned on the top of figures. Additional summary statistics are provided in Tables 8–9. All statistical test outcomes associated with this figure are provided in Table 10*.

**Table 10:** Results of statistical tests performed for physiological properties between different blades of the dentate gyrus (SPB: Suprapramidal blade, IPB: Infrapyramidal blade; Figs. 6–8). *p* values are provided for the Wilcoxon Rank sum test. Statistically significant *p* values are represented in boldface font.

| Measurement, unit | Symbol | Dorsal<br>SPB vs. IPB | Intermediate<br>SPB vs. IPB | Ventral<br>SPB vs. IPB |
| --- | --- | --- | --- | --- |
| *Subthreshold measurements* |  |  |  |  |
| Resting membrane potential, mV | $V_{\text{RMP}}$ | **0.01** | 0.36 | 0.63 |
| Input resistance, MΩ | $R_{\text{in}}$ | 0.06 | 0.52 | 0.11 |
| Temporal summation ratio | $S_{\alpha}$ | 0.22 | **0.03** | 0.67 |
| Sag ratio, % | $Sag$ | 0.68 | 0.63 | 0.09 |
| Resonance frequency, Hz | $f_R$ | 0.60 | 0.38 | 0.079 |
| Maximum impedance amplitude, MΩ | $\vert Z\vert _{\text{max}}$ | 0.59 | 0.27 | 0.41 |
| Resonance strength | $Q$ | 0.12 | 0.51 | 0.37 |
| Total inductive phase, rad·Hz | $\Phi_L$ | 0.27 | 0.96 | 0.53 |
| *Suprathreshold measurements* |  |  |  |  |
| Firing rate at 50 pA, Hz | $f_{50}$ | — | 0.97 | 0.120 |
| Firing rate at 100 pA, Hz | $f_{100}$ | 0.80 | **0.02** | **0.006** |
| Firing rate at 150 pA, Hz | $f_{150}$ | 0.95 | 0.19 | **0.001** |
| Firing rate at 200 pA, Hz | $f_{200}$ | 0.46 | 0.31 | **0.002** |
| Firing rate at 250 pA, Hz | $f_{250}$ | 0.05 | 0.75 | **0.0009** |
| Firing rate at 500 pA, Hz | $f_{500}$ | 0.09 | 0.35 | **0.016** |
| Firing rate at 750 pA, Hz | $f_{750}$ | 0.15 | 0.06 | 0.05 |
| Firing rate at 1000 pA, Hz | $f_{1000}$ | 0.45 | **0.002** | 0.37 |
| Firing rate at 1250 pA, Hz | $f_{1250}$ | 0.70 | **0.03** | 0.62 |
| Peak action potential voltage, mV | $V_{\text{peak}}$ | **0.0013** | 0.88 | 0.21 |
| Action potential amplitude, mV | $V_{\text{AP}}$ | 0.1000 | 0.30 | 0.20 |
| Action potential threshold, mV | $V_{\text{th}}$ | 0.552 | 0.19 | 0.19 |
| Action potential halfwidth, ms | $T_{\text{APHW}}$ | **0.03** | 0.05 | 0.84 |
| Peak dV/dt, V/s | $dV/dt\vert _{\text{max}}$ | **0.01** | 0.35 | 0.25 |
| Minimum dV/dt, V/s | $dV/dt\vert _{\text{min}}$ | **0.02** | 0.58 | 0.33 |
| Latency to first spike, ms | $T_{1\text{AP}}$ | **0.04** | **0.04** | **0.01** |
| First inter-spike interval, ms | $T_{1\text{ISI}}$ | 0.17 | **0.011** | **0.02** |

In the dorsal dentate gyrus, although the resting membrane potential of IPB granule cells was significantly hyperpolarized compared to their SPB counterparts (Fig. 6*B*), we did not find significant differences in their subthreshold (Fig. 6*B–C*) or suprathreshold (Fig. 6*D–E*) excitability properties. A significant reduction in action potential peak voltage without significant changes to action potential amplitude (Fig. 6*F*) was broadly consistent with the hyperpolarized RMP (Fig. 6*B*). We also found small, yet significant differences in several action potential properties of SPB *vs*. IPB granule cells (Fig. 6*G*).

For granule cells from the intermediate section, consistent with previous observations from the blades of intermediate hippocampus (Mishra & Narayanan, 2020a), we found most subthreshold and suprathreshold measurements to be comparable between the SPB and IPB (Fig. 7). The firing frequency of SPB granule cells was higher than their IPB counterparts for higher current injections (Fig. 7*D–E*). We found small, yet significant differences in latency to first spike and in first interspike interval (ISI) between SPB and IPB granule cells (Fig. 7*G*).

In striking contrast to the absence of significant blade differences in excitability properties of dorsal and intermediate granule cells, we found significant differences between firing rates of granule cells in SPB *vs*. IPB in ventral DG (Fig. 8). Specifically, despite the absence of significant differences in resting membrane potential or subthreshold properties (Fig. 8*B–C*), we found significant reductions in the action potential firing frequency of granule cells in the ventral SPB compared to those in ventral IPB (Fig. 8*D–E*). Consistent with higher suprathreshold excitability of IPB granule cells, we found them to switch to depolarization-induced block with relatively lower current injections compared to their SPB counterparts (Fig. 8*D–E*). Blade-specific differences in action potential properties, especially significant reductions of both latency to first spike and first ISI (Fig. 8*G*), were consistent with higher firing rates of IPB granule cells compared to SPB granule cells. Although not significantly different, the distribution of AP threshold showed a downward shift and that of peak *dV*/*dt* indicated an upward shift in IPB cells, both of which are consistent with higher firing rates of IPB cells (Fig. 8*G*).

Together, our analyses showed that ventral granule cells in the infrapyramidal blade manifested higher firing rates compared to their counterparts in the suprapyramidal blade, without significant differences in subthreshold excitability. These blade-specific differences were limited to the ventral DG, with negligible distinctions between most physiological properties of GCs in the two blades of either dorsal or intermediate DG. The distribution of the different physiological measurements (Fig. 6–8) and various measures of heterogeneity (Tables 4–9) emphasized heterogeneities within each of the two blades of GCs across the dorsoventral axis.

### Blade-specific contributions to dorsoventral gradient in granule cell excitability

We had found a gradient in subthreshold (Fig. 2) and suprathreshold (Fig. 4) excitability of DG granule cells along the dorsal-to-ventral axis. In addition, within the ventral dentate gyrus, the IPB granule cells manifested higher suprathreshold excitability without significant changes to subthreshold excitability (Fig. 8). How do SPB and IPB granule cells across the dorsoventral compare with each other? Did the gradient in excitability across the dorsoventral axis maintain individually along both SPB and IPB or was it confined to only one of them? How much of the dorsal-to-ventral gradient in excitability was contributed by blade-specific differences in the ventral DG?

In addressing these questions, we assessed the electrophysiological properties of granule cells in six categories depending on their location (Figs. 9–10; Tables 10–11): dorsal SPB (*n* = 23), dorsal IPB (*n* = 26), intermediate SPB (*n* = 21), intermediate IPB (*n* = 21), ventral SPB (*n* = 20), and ventral IPB (*n* = 23). Individually comparing subthreshold properties of SPB and IPB GCs across the dorsoventral axis, we found a small yet significantly depolarized RMP of ventral IPB cells as compared to their dorsal counterparts (Fig. 9*B*). The increase in input resistance and maximal impedance amplitude along the dorsal-to-ventral axis was maintained within both SPB and IPB as well (Fig. 9*B*). The other subthreshold measurements did not show significant changes across the dorsoventral axis within either SPB or IPB (Fig. 9*B*). Together, these analyses confirmed that the dorsal-to-ventral gradient in subthreshold excitability was observable both in SPB and IPB (Fig. 9), together contributing to the overall dorsoventral gradient in subthreshold excitability of granule cells observed earlier (Fig. 2).

**Figure 9.**
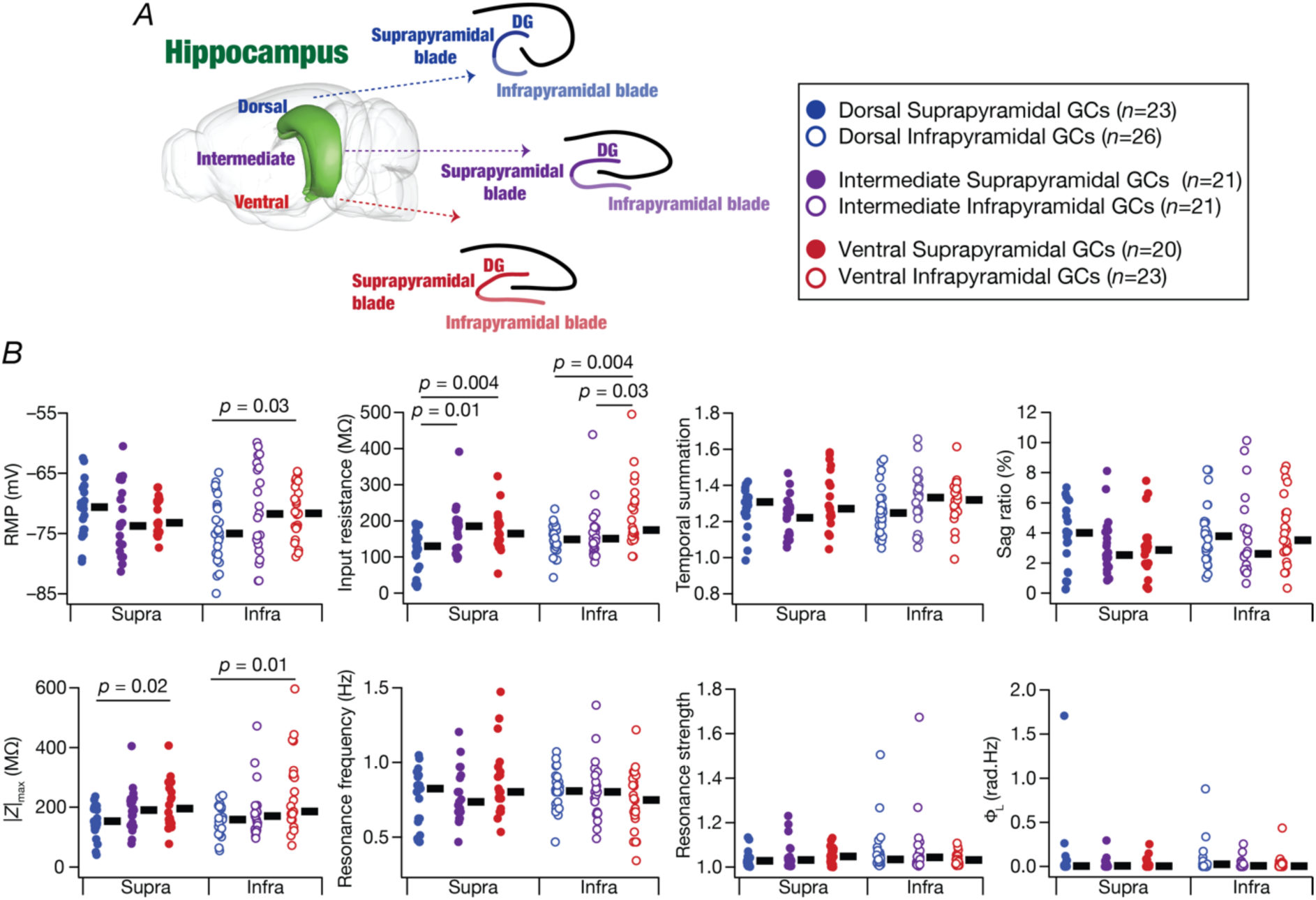
Analyses of blade-wise contributions to the dorsoventral subthreshold excitability gradient in dentate gyrus granule cells. *A*, Schematic of rodent brain showing the anatomical structure of transverse section of dorsal, intermediate and ventral hippocampus (Image from Scalable Brain Atlas (Bakker *et al*., 2015)). *B*, Beeswarm plots depicting subthreshold measurements from granule cells of dorsal, intermediate, and ventral dentate gyrus: resting membrane potential (RMP), *V*_*RMP*_ (mV), input resistance, *R*_*in*_ (MΩ), temporal summation, *S*_*⍺*_, Sag (%), impedance amplitude, |*Z*|_*max*_ (MΩ), resonance frequency, *f*_*R*_ (Hz), resonance strength, *Q*, and total inductive phase, Φ_L_(rad.Hz) from all the recorded granule cells from suprapyramidal and infrapyramidal blades of dorsal, intermediate, and ventral dentate gyrus. The thick lines on right of each beeswarm plot represent the median for the specified population. Dorsal suprapyramidal GCs, *n* = 23; dorsal infrapyramidal GCs, *n* = 26; intermediate suprapyramidal GCs, *n* = 21; intermediate infrapyramidal GCs, *n* = 21; ventral suprapyramidal GCs, *n* = 20; ventral infrapyramidal GCs, *n* = 23. *Statistical test: Wilcoxon rank sum test was performed between two groups, significant p values are mentioned on the top of figures. Additional summary statistics are provided in Tables 4–9. All statistical test outcomes associated with this figure are provided in Tables 11–12*.

**Figure 10.**
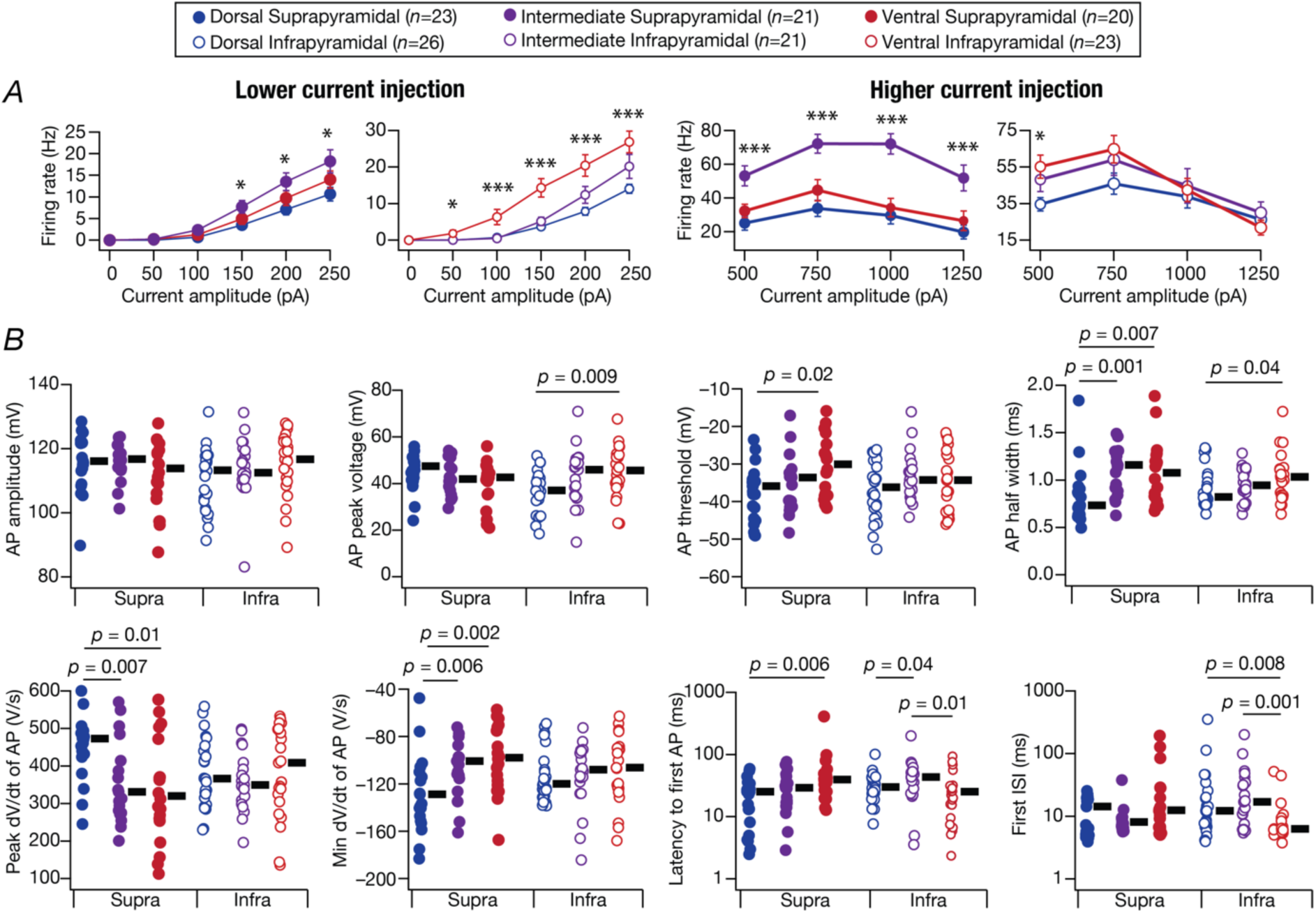
Analyses of blade-wise contributions to the dorsoventral suprathreshold excitability gradient in dentate gyrus granule cells. *A*, *f* − *I* curves for the different neuron categories for lower (left) and higher (right) current injection protocols. *p values: * <0.05, ** < 0.005, *** < 0.0005*. *B*, Beeswarm plots comparing the action potential measurements in all the 6 categories of neurons. The thick lines on right of each beeswarm plot represent the median for the specified population. Dorsal suprapyramidal GCs, *n* = 23; dorsal infrapyramidal GCs, *n* = 26; intermediate suprapyramidal GCs, *n* = 21; intermediate infrapyramidal GCs, *n* = 21; ventral suprapyramidal GCs, *n* = 20; ventral infrapyramidal GCs, *n* = 23. *Statistical test: Wilcoxon rank sum test was performed between two groups, significant p values are mentioned on the top of figures. Additional summary statistics are provided in Tables 4–9. All statistical test outcomes associated with this figure are provided in Tables 11–12*.

**Table 11:** Results of statistical tests performed for physiological properties between different dorsoventral suprapyramidal blade (SPB) groups (D: Dorsal, I: Intermediate, V: Ventral; Figs. 9–10). Statistically significant *p* values are represented in boldface font.

| Measurement, unit | Symbol | KW test | Pairwise Wilcoxon Rank sum test |  |  |
| --- | --- | --- | --- | --- | --- |
| | | | $V$ vs. $I$ | $V$ vs. $D$ | $I$ vs. $D$ |
| Subthreshold measurements |  |  |  |  |  |
| Resting membrane potential, mV | $V_{\text{RMP}}$ | 0.37 | 0.41 | 0.41 | 0.21 |
| Input resistance, MΩ | $R_{in}$ | **0.006** | 0.78 | **0.004** | **0.01** |
| Temporal summation ratio | $S_{\alpha}$ | 0.13 | 0.11 | 0.86 | 0.06 |
| Sag ratio, % | $Sag$ | 0.69 | 0.95 | 0.41 | 0.53 |
| Resonance frequency, Hz | $f_R$ | 0.35 | 0.14 | 0.32 | 0.95 |
| Maximum impedance amplitude, MΩ | $\vert Z\vert _{max}$ | **0.04** | 0.49 | **0.02** | 0.08 |
| Resonance strength | $Q$ | 0.42 | 0.72 | 0.21 | 0.37 |
| Total inductive phase, rad·Hz | $\Phi_L$ | 0.85 | 0.66 | 1 | 0.61 |
| Suprathreshold measurements |  |  |  |  |  |
| Firing rate at 50 pA, Hz | $f_{50}$ | 0.55 | 1 | 0.29 | 0.31 |
| Firing rate at 100 pA, Hz | $f_{100}$ | 0.19 | 0.14 | 0.55 | 0.14 |
| Firing rate at 150 pA, Hz | $f_{150}$ | 0.08 | 0.12 | 0.66 | **0.02** |
| Firing rate at 200 pA, Hz | $f_{200}$ | **0.04** | 0.16 | 0.27 | **0.01** |
| Firing rate at 250 pA, Hz | $f_{250}$ | 0.06 | 0.17 | 0.24 | **0.02** |
| Firing rate at 500 pA, Hz | $f_{500}$ | **0.002** | **0.01** | 0.18 | **0.001** |
| Firing rate at 750 pA, Hz | $f_{750}$ | **$8.8 \times 10^{-5}$** | **0.003** | 0.2 | **$4.7 \times 10^{-5}$** |
| Firing rate at 1000 pA, Hz | $f_{1000}$ | **$4.9 \times 10^{-5}$** | **$4 \times 10^{-4}$** | 0.61 | **$8.3 \times 10^{-5}$** |
| Firing rate at 1250 pA, Hz | $f_{1250}$ | **0.001** | **0.01** | 0.45 | **$6 \times 10^{-4}$** |
| Peak action potential voltage, mV | $V_{\text{peak}}$ | 0.22 | 0.73 | 0.07 | 0.31 |
| Action potential amplitude, mV | $V_{AP}$ | 0.48 | 0.27 | 0.36 | 0.81 |
| Action potential threshold, mV | $V_{th}$ | **0.036** | 0.06 | **0.02** | 0.29 |
| Action potential halfwidth, ms | $T_{\text{APHW}}$ | **0.001** | 0.58 | **0.007** | **0.001** |
| Peak dV/dt, V/s | $dV/dt\vert _{\text{max}}$ | **0.01** | 0.61 | **0.01** | **0.007** |
| Minimum dV/dt, V/s | $dV/dt\vert _{\text{min}}$ | **0.002** | 0.38 | **0.002** | **0.006** |
| Latency to first spike, ms | $T_{1\text{AP}}$ | **0.01** | 0.08 | **0.006** | 0.26 |
| First inter-spike interval, ms | $T_{1\text{ISI}}$ | 0.18 | 0.1 | 0.12 | 0.73 |

**Table 12:** Results of statistical tests performed for physiological properties between different dorsoventral infrapyramidal blade (IPB) groups (D: Dorsal, I: Intermediate, V: Ventral; Figs. 9–10). Statistically significant *p* values are represented in boldface font.

| Measurement, unit | Symbol | KW test | Pairwise Wilcoxon Rank sum test |  |  |
| --- | --- | --- | --- | --- | --- |
|  |  |  | *V* vs. *I* | *V* vs. *D* | *I* vs. *D* |
| Subthreshold measurements |  |  |  |  |  |
| Resting membrane potential, mV | $V_{RMP}$ | 0.1 | 0.87 | **0.03** | 0.15 |
| Input resistance, MΩ | $R_{in}$ | **0.013** | **0.03** | **0.004** | 0.79 |
| Temporal summation ratio | $S_{\alpha}$ | 0.13 | 0.83 | 0.06 | 0.13 |
| Sag ratio, % | $Sag$ | 0.5 | 0.34 | 0.96 | 0.28 |
| Resonance frequency, Hz | $f_R$ | 0.32 | 0.24 | 0.17 | 0.8 |
| Maximum impedance amplitude, MΩ | $\vert Z\vert _{max}$ | 0.05 | 0.09 | **0.017** | 0.71 |
| Resonance strength | $Q$ | 0.58 | 0.45 | 0.31 | 0.97 |
| Total inductive phase, rad·Hz | $\Phi_L$ | 0.61 | 0.96 | 0.33 | 0.5 |
| Suprathreshold measurements |  |  |  |  |  |
| Firing rate at 50 pA, Hz | $f_{50}$ | **0.015** | 0.09 | **0.01** | 0.28 |
| Firing rate at 100 pA, Hz | $f_{100}$ | $4 \times 10^{-4}$ | $6 \times 10^{-6}$ | **0.003** | 0.21 |
| Firing rate at 150 pA, Hz | $f_{150}$ | $2.3 \times 10^{-5}$ | $4 \times 10^{-4}$ | $2 \times 10^{-5}$ | 0.35 |
| Firing rate at 200 pA, Hz | $f_{200}$ | $1.6 \times 10^{-4}$ | **0.006** | $5 \times 10^{-5}$ | 0.19 |
| Firing rate at 250 pA, Hz | $f_{250}$ | $6 \times 10^{-4}$ | **0.02** | $1 \times 10^{-4}$ | 0.2 |
| Firing rate at 500 pA, Hz | $f_{500}$ | **0.04** | 0.47 | **0.015** | 0.11 |
| Firing rate at 750 pA, Hz | $f_{750}$ | 0.2 | 0.36 | 0.09 | 0.36 |
| Firing rate at 1000 pA, Hz | $f_{1000}$ | 0.84 | 0.92 | 0.6 | 0.67 |
| Firing rate at 1250 pA, Hz | $f_{1250}$ | 0.61 | 0.37 | 0.71 | 0.47 |
| Peak action potential voltage, mV | $V_{peak}$ | **0.02** | 0.49 | **0.009** | 0.08 |
| Action potential amplitude, mV | $V_{AP}$ | 0.13 | 0.39 | 0.06 | 0.21 |
| Action potential threshold, mV | $V_{th}$ | 0.3 | 0.49 | 0.38 | 0.14 |
| Action potential halfwidth, ms | $T_{APHW}$ | 0.12 | 0.43 | **0.04** | 0.23 |
| Peak dV/dt, V/s | $dV/dt\vert _{max}$ | 0.77 | 0.47 | 0.77 | 0.69 |
| Minimum dV/dt, V/s | $dV/dt\vert _{min}$ | 0.37 | 0.67 | 0.25 | 0.23 |
| Latency to first spike, ms | $T_{1AP}$ | **0.017** | **0.019** | 0.07 | **0.049** |
| First inter-spike interval, ms | $T_{1ISI}$ | **0.002** | **0.001** | **0.008** | 0.4 |

Turning to blade-wise analyses of suprathreshold excitability along the dorsoventral axis, we found significant contributions from the IPB in the enhanced firing frequency of ventral GCs, for current injections in the range of 50–250 pA (Fig. 10*A*; Table 11; see panel E of Figs. 6–8 for the distribution of firing rates for all current values). The higher excitability of ventral IPBs was consistent with a reduction in their average firing rates at higher current injections (above 250 pA) owing to the onset of depolarization-induced block. Thus, the dorsal-to-ventral gradient in firing rate in IPB granule cells (Fig. 10*A*) was consistent with the overall trend observed earlier for the pooled data across both blades (Fig. 4). In contrast, the firing rates of granule cells from the intermediate hippocampus were the highest for the SPB across all measured current injections (Fig. 10*A*; Table 12; see panel E of Figs. 6–8 for the distribution of firing rates for all current values). Together, whereas ventral granule cells showed the highest firing rates among IPB neurons, intermediate granule cells were highly excitable among SPB neurons. Dorsal granule cells showed the lowest firing rate excitability irrespective of which blade they belonged to.

Among action potential measurements, there was a significant increase in the AP peak voltage between dorsal and ventral IPB neurons, with no significant differences in AP amplitude (Fig. 10*B*). Action potential threshold was significantly depolarized in SPB dorsal neurons compared to their ventral counterparts (Fig. 10*B*). Consistent with our observations on the pooled population (Fig. 4), measurements associated with the AP repolarization kinetics showed gradients in individual blades as well (Fig. 10*B*). Specifically, both AP halfwidth and minimum *dV*/*dt* of granule cells increased along the dorsal-to-ventral axis for both SPB and IPB, although min *dV*/*dt* of IPB granule cells was not significantly different across the three groups (Fig. 10*B*; Tables 11–12). The repolarization kinetics in dorsal SPB granule cells were the fastest with significantly fast repolarization *dV*/*dt* and significantly thinner action potentials (Fig. 10*B*; Tables 11–12). Among SPB granule cells, peak *dV*/*dt* was significantly higher for dorsal cells, with no differences among IPB granule cells (Fig. 10*B*; Tables 11–12). We also found small yet significant changes along the dorsoventral axis in latency to first AP in both blades and in first ISI within IPB but not SPB (Fig. 10*B*; Tables 11–12). Together, among AP measurements, the fast repolarization kinetics of dorsal granule cells was consistent across both SPB and IPB, while other measurements showed distinct dorsoventral differences between SPB and IPB.

Together, with specific reference to dorsoventral measurements from the two DG blades, the dorsal-to-ventral gradients in subthreshold excitability measurements were consistent across IPB and SPB. However, with reference to suprathreshold excitability, the consistent observation was that the dorsal neurons showed the lowest action potential firing frequency along the dorsoventral axis. Whereas the ventral neurons showed the highest firing frequency among IPB granule cells, the intermediate neurons were the most excitable among SPB granule cells. Finally, the dorsal-to-ventral gradient in repolarization kinetics was consistent between both SPB and IPB, with dorsal granule cells showing the thinnest action potentials.

## DISCUSSION

We systematically characterized the intrinsic electrophysiological properties of dentate gyrus (DG) granule cells (GCs) along the dorsoventral axis of the hippocampus and uncovered a robust gradient in cellular excitability. Specifically, we observed a progressive gradient in subthreshold excitability measures, in firing rate, and in repolarization kinetics of action potentials along the dorsal-to-ventral axis of the hippocampus. Our measurements also demonstrated that granule cells acted as class I integrators that lacked strong resonance properties across the dorsoventral axis. We performed pairwise correlation and dimensionality reduction analyses to reveal weak dependencies across physiological measurements and the absence of distinct clusters for dorsal, intermediate, or ventral cells. Together, these observations emphasized the manifestation of widespread heterogeneity that needs to be accounted for in all analyses.

A more granular analysis of these measurements revealed that this gradient in excitability was not homogeneous across the two blades of the DG, but was shaped by blade-specific contributions from the infrapyramidal blade (IPB) and suprapyramidal blade (SPB). Specifically, we found ventral IPB granule cells to manifest higher firing rates compared to ventral SPB granule cells, without significant differences in subthreshold excitability. Importantly, these blade-specific differences were limited to the ventral DG. The dorsal-to-ventral gradient showing an increase in subthreshold excitability measurements was consistent across both IPB and SPB. However, while dorsal neurons showed the lowest action potential firing frequency irrespective of which blade they belonged to, ventral neurons showed the highest firing frequency among IPB granule cells but intermediate neurons were the highest excitable among SPB granule cells. The dorsal-to-ventral gradient in repolarization kinetics was consistent between both SPB and IPB, with dorsal granule cells showing the thinnest action potentials. Together, our findings provide evidence for a dorsal-to-ventral gradient in GC intrinsic excitability, highlighting both a progressive increase in excitability along the dorsal-to-ventral axis and a blade-specific granularity of firing properties.

### Dorsoventral differences in the hippocampal formation

The rodent hippocampus is a heterogeneous structure with neurons in different subregions showing very different properties (Fanselow & Dong, 2010; Strange *et al*., 2014; Cembrowski *et al*., 2016; Cembrowski & Spruston, 2019). These heterogeneities span different axes, with well-studied differences between dorsal *vs*. ventral, proximal *vs.* distal, and deep *vs.* superficial hippocampal neurons (Jarsky *et al*., 2008; Thompson *et al*., 2008; Valero *et al*., 2015; Maroso *et al*., 2016; Sun *et al*., 2017; Valero & de la Prida, 2018). Despite this broader context, the dentate gyrus has received relatively limited attention in terms of dorsoventral electrophysiological profiling of neuronal intrinsic properties. Our study therefore provides a critical missing link, showing that, like CA1 pyramidal and entorhinal neurons, DG granule cells also exhibit a systematic dorsoventral gradient in intrinsic excitability.

Our findings align with and extend prior studies demonstrating dorsoventral gradients in intrinsic excitability across the hippocampal formation. In CA1 pyramidal neurons, several studies have reported that ventral cells are more excitable than dorsal cells, characterized by higher input resistance and increased firing rates (Dougherty *et al*., 2012; Marcelin *et al*., 2012; Malik *et al*., 2016). Similar gradients have also been described in medial entorhinal cortex (MEC) stellate cells, where ventral neurons exhibit slower membrane time constants, lower resonance frequency, and enhanced synaptic integration (Giocomo *et al*., 2007; Garden *et al*., 2008; Giocomo & Hasselmo, 2008; Pastoll *et al*., 2020). These observations collectively suggest that intrinsic membrane properties are regionally tuned to support distinct computational demands along the hippocampal axis. The dorsal-to-ventral increase in subthreshold and suprathreshold excitability in DG granule cells reported here is broadly consistent with the direction of such gradients in the CA1 pyramidal neurons and in entorhinal stellate cells.

### Regional heterogeneities within the DG

By dissecting the relative contributions of the infrapyramidal and suprapyramidal blades across regions, we add a new layer of spatial resolution to the dorsoventral gradient in intrinsic properties of granule cells. Our analyses reveal that dorsoventral differences in intrinsic properties are further modulated by intra-DG blade-specific heterogeneities. Our findings add to the emerging view that region- and blade-specific tuning of intrinsic membrane properties is a widespread organizational principle in the hippocampal formation, likely reflecting adaptation to distinct input-output functions along the longitudinal axis.

There are several well-established differences across these two blades (Amaral *et al*., 2007), including morphological differences (Desmond & Levy, 1982, 1985; Green & Juraska, 1985; Claiborne *et al*., 1990; Schneider *et al*., 2014; Gallitano *et al*., 2016), intrinsic properties and synaptic plasticity profiles (Mishra & Narayanan, 2020a; Strauch *et al*., 2025), connectivity patterns (Claiborne *et al*., 1986), the ratio of basket cells to granule cells (Seress & Pokorny, 1981), and activity levels (Chawla *et al*., 2005; Ramirez-Amaya *et al*., 2005; Ramirez-Amaya *et al*., 2006; Marrone *et al*., 2012a; Marrone *et al*., 2012b; Satvat *et al*., 2012; Ramirez-Amaya *et al*., 2013). Our observations from the intermediate hippocampus are consistent with our previous study (Mishra & Narayanan, 2020a), where intrinsic properties were heterogenous but comparable across the blades. Those recordings were performed with spontaneous synaptic activity intact, in the absence of synaptic blockers. Our recordings in the presence of synaptic blockers from the intermediate hippocampus find comparable measurements and heterogeneities of intrinsic physiological properties across the two blades of the DG.

We extended our analyses to granule cells in DG blades across the dorsoventral axis revealing strong blade-specific differences in the ventral DG, but with largely comparable properties across the blades of the intermediate and dorsal DG. These blade-specific excitability profiles may align with distinct connectivity patterns across blades, and may participate in parallel information streams related to spatial and emotional processing (Amaral *et al*., 2007). Such blade-specific tuning of intrinsic excitability may therefore serve to filter or amplify distinct categories of input along the longitudinal axis, supporting the DG’s proposed role as a context-sensitive gate for information entering the hippocampus.

### Mechanistic origins of excitability gradients

Our quantitative assessment of the several electrophysiological properties across the dorsoventral axis provide pointers to potential mechanistic origins behind the observed excitability gradients. First, with reference to subthreshold gradients, the higher input resistance and maximum impedance amplitude observed in ventral granule cells (Fig. 2) point to potential gradients in morphological properties or molecular composition (including ion channel expression and other transmembrane or cytoplasmic components). While morphology is a strong modulator of excitability (Mainen & Sejnowski, 1996; Narayanan *et al*., 2003; Dhupia *et al*., 2015), the gradient in subthreshold excitability could also be mediated by gradients in the several resting conductances. Specifically, excitability gradients could be mediated by expression profiles of leak channels, Ba-sensitive inward-rectifying potassium (K_ir_) channels, or hyperpolarization-activated cyclic-nucleotide gated (HCN) nonspecific cation channels, each of which are known to express in DG granule cells (Stegen *et al*., 2009; Stegen *et al*., 2012; Surges *et al*., 2012; Mishra & Narayanan, 2021a; Mishra & Narayanan, 2022). Irrespective of the location of the granule cells within the dorsoventral axis, we observed progressive hyperpolarization-induced reductions in input resistance (Fig. 2*I*) and maximal impedance amplitude (Fig. 2*L*), accompanied by a hyperpolarization-induced increase in sag (Fig. 2*J*). These observations point to the putative expression of hyperpolarization-activated channels such as HCN and K_ir_ in granule cells across the dorsoventral axis. The absence of strong changes in RMP (Fig. 2*A*) suggest that ion-channel gradients, if present, could be in both HCN and K_ir_ channels. While enhanced HCN and K_ir_ channels expressions affect input resistance similarly, their effects on RMP are opposite (Mishra & Narayanan, 2021a; Mishra & Narayanan, 2022), together predicting a joint gradient of reduction in HCN and K_ir_ channel expression along the dorsal-to-ventral axis. Across the dorsoventral axis of the DG, our measurements don’t show significant differences in subthreshold excitability between SPB and IPB (Figs. 6–8), suggesting such gradients in morphological or resting conductances to be spanning the dorsoventral axis and not extending across blades.

Suprathreshold excitability gradient in AP firing rates across the dorsoventral axis (Fig. 4*B*) could be an extension of the subthreshold gradients in input resistance. In addition, strong changes in repolarization kinetics across the dorsoventral axis, manifesting as increasing AP half-width (Fig. 4*F*) as well as reducing rate of repolarization (Fig. 4*H*), point to potential gradients in potassium channels mediating AP repolarization. Specifically, one or more of *A*-type, *D*-type, big-conductance calcium-activated potassium (BK), or delayed rectifier potassium channels could be showing reduced activity along the dorsal-to-ventral axis towards implementing slower repolarization kinetics. Although the absence of strong changes in AP amplitude (Fig. 4*C–D*) point to minimal changes to voltage-gated sodium (Na_v_) channels, there are significant differences in AP threshold (Fig. 4*E*) and peak *dV*/*dt* (Fig. 4*G*) which could originate from changes in Na_v_ channel expression gradients. The observed blade-specific differences in granule cell firing properties in the absence of changes in subthreshold excitability, especially in the ventral hippocampus (Fig. 8), are likely due to differences in ion channel expression profiles and/or properties. Specifically, ventral IPB granule cells elicit more action potentials compared to the SPB counterparts (Fig. 8*D–E*), with significant reductions in both latency to first spike and first ISI (Fig. 8*G*). These changes point to potential blade-specific differences in one or more of Na_v_ channels, *T*-type calcium channels, *A-*type potassium, or *D*-type potassium channels.

Importantly, the mapping between ion-channels and cellular measurements in granule cells has been shown to be many-to-many (Beining *et al*., 2017; Mishra & Narayanan, 2019, 2021a; Kumari & Narayanan, 2024). Thus, dorsoventral gradients and blade-specific differences in several channels (and other components, including morphological and molecular) could collectively contribute to the observed differences in physiological measurements (Rathour & Narayanan, 2019; Albantakis *et al*., 2024). A systematic blade-specific assessment of different ion channels from granule cells across the dorsoventral axis using molecular profiling methods, cell-attached recordings, pharmacological blockers, and computational modeling should therefore be performed to elucidate the mechanistic origins of heterogeneities and gradients in physiological properties.

### Physiological and pathological implications for within-cell-type heterogeneities

Heterogeneities spanning morphological characteristics, molecular composition, and physiological properties within a given neuronal subtype are ubiquitous (Cembrowski & Spruston, 2019; Mishra & Narayanan, 2020a; Moradi Chameh *et al*., 2021; Peng *et al*., 2021; Scala *et al*., 2021; Mittal & Narayanan, 2022; Han *et al*., 2023; Yao *et al*., 2023; Dahmen *et al*., 2025; Planert *et al*., 2025). While the dorsoventral and blade differences are crucial observations, our recordings also unveil widespread heterogeneity in intrinsic physiology of hippocampal granule cells, even within a given dorsal-to-ventral section and even when restricted to a specific blade (Figs. 6–10; Tables 4–9). These within-type heterogeneities have been shown to play beneficial or detrimental roles in neural circuit function, depending on several factors including the circuit under consideration, the intrinsic properties of neurons therein, the specific connectivity pattern, the task implemented by the network, the form and degree of the heterogeneity, and interactions among different forms of heterogeneities (Wang & Buzsaki, 1996; Brunel & Hakim, 1999; Tikidji-Hamburyan *et al*., 2015; Mishra & Narayanan, 2016, 2021b; Mittal & Narayanan, 2021; Gast *et al*., 2024; Dahmen *et al*., 2025; Saini & Narayanan, 2025). With specific reference to the dentate gyrus, the manifestation of heterogeneities has been shown to promote resilience and robustness of channel decorrelation and pattern separation (Mishra & Narayanan, 2019, 2021b; Saini & Narayanan, 2025). Future studies could explore the specific impact of dorsoventral gradients and blade-specific differences reported here on pattern separation and channel decorrelation.

The intrinsic excitability gradient we observed along the dorsoventral axis likely has important consequences for the established functional roles of DG in cognition and behavior, especially considering the dichotomy of roles that have been attributed to dorsal *vs*. ventral hippocampus (Pothuizen *et al*., 2004; Fanselow & Dong, 2010; Kheirbek *et al*., 2013; Weeden *et al*., 2014; Kempadoo *et al*., 2016; Kirk *et al*., 2017; Kesner, 2018; Woods *et al*., 2020; Yeates *et al*., 2020; Dirven *et al*., 2022). Apart from assessing the roles of this dorsoventral gradient in intrinsic excitability to novelty detection, engram formation, exploratory drive, contextual fear conditioning, pattern separation, reward encoding, emotional salience, and social recognition, studies should also assess if there are dorsoventral gradients in place-field properties of granule cells (Danielson *et al*., 2017; GoodSmith *et al*., 2017; Senzai & Buzsaki, 2017; Zhang & Jonas, 2020; GoodSmith *et al*., 2022), similar to those found in other parts of the hippocampal formation (Hafting *et al*., 2005; Kjelstrup *et al*., 2008; Royer *et al*., 2010). Importantly, gradients in intrinsic properties of the principal cells is one piece of the puzzle of how they contribute to different aspects of active behavior. Future studies should systematically explore dorsoventral gradients in neuronal morphology, interneuronal and astrocytic intrinsic properties, synaptic/intrinsic plasticity profiles, synaptic and neuron-glia interactions. Furthermore, as synaptic and intrinsic properties could be altered in a behavioral-state dependent manner by different neuromodulatory inputs that arrive on to the dentate gyrus, it is essential to assess neuronal intrinsic properties *in vivo* under different behavioral conditions. Such approaches could explore putative links between intrinsic excitability and behaviorally relevant neuronal recruitment, offering a deeper understanding of how intrinsic properties shape firing output and stable continual learning under naturalistic input conditions (Park *et al*., 2016; Titley *et al*., 2017; Josselyn & Frankland, 2018; Rao-Ruiz *et al*., 2019; Josselyn & Tonegawa, 2020; Lau *et al*., 2020; Mishra & Narayanan, 2021c). Together, these directions will deepen our understanding of how spatially organized cellular properties contribute to the rich functional heterogeneity of the hippocampus.

Finally, these regionally tuned excitability profiles could also have implications for neurological and psychiatric conditions that differentially affect the hippocampus along the dorsoventral axis (Scharfman *et al*., 2000; Scharfman *et al*., 2002; Scharfman, 2007; Sutula & Dudek, 2007; Winner *et al*., 2011; Hagihara *et al*., 2013; Patricio *et al*., 2013; Yu *et al*., 2014; Cho *et al*., 2015; Leal & Yassa, 2015; Arnold *et al*., 2019; Botterill *et al*., 2019; Stober *et al*., 2023). For example, our findings of elevated excitability of ventral GCs could contribute to an increased vulnerability of these neurons to pathological hyperexcitability or maladaptive plasticity in different neurological disorders. On the other hand, impaired recruitment of granule cells due to hypoexcitability or loss of heterogeneities among granule cells could degrade pattern separation and contribute to memory deficits observed in different neurological disorders (Yassa & Stark, 2011; Rich *et al*., 2022). Together, understanding how intrinsic excitability is spatially organized would help in clarifying the circuit-level mechanisms underlying both healthy cognition and diverse disease states where the DG has been strongly implicated.

## Author contributions

S.K. and R.N. designed experiments; S.K. performed experiments; S.K. analyzed data; S.K. and R.N. wrote the paper.

## Data availability statement

All data and associated analyses outcomes required for assessment of this study are available as part of the manuscript.

## Acknowledgments

The authors thank members of the cellular neurophysiology laboratory for helpful discussions and for comments on a draft of this manuscript.

